# Ultra-sensitive isotope probing to quantify activity and substrate assimilation in microbiomes

**DOI:** 10.1101/2021.03.29.437612

**Authors:** Manuel Kleiner, Angela Kouris, Marlene Jensen, Grace D’Angelo, Yihua Liu, Abigail Korenek, Nikola Tolić, Timo Sachsenberg, Janine McCalder, Mary S. Lipton, Marc Strous

## Abstract

Stable isotope probing (SIP) approaches are a critical tool in microbiome research to determine associations between species and substrates. The application of these approaches ranges from studying microbial communities important for global biogeochemical cycling to host-microbiota interactions in the intestinal tract. Current SIP approaches, such as DNA-SIP or nanoSIMS, are limited in terms of sensitivity, resolution or throughput. Here we present an ultra-sensitive, high-throughput protein-based stable isotope probing approach (Protein-SIP), which cuts cost for labeled substrates by ∼90% as compared to other SIP and Protein-SIP approaches and thus enables isotope labeling experiments on much larger scales and with higher replication. It allows for the determination of isotope incorporation into microbiome members with species level resolution using standard metaproteomics LC-MS/MS measurements. The analysis has been implemented as an open-source application (https://sourceforge.net/projects/calis-p/). We demonstrate sensitivity, precision and accuracy using bacterial cultures and mock communities with different labeling schemes. Furthermore, we benchmark our approach against two existing Protein-SIP approaches and show that in the low labeling range used our approach is the most sensitive and accurate. Finally, we measure translational activity using 18O heavy water labeling in a 63-species community derived from human fecal samples grown on media simulating two different diets. Activity could be quantified on average for 27 species per sample, with 9 species showing significantly higher activity on a high protein diet, as compared to a high fiber diet. Surprisingly, among the species with increased activity on high protein were several *Bacteroides* species known as fiber consumers. Apparently, protein supply is a critical consideration when assessing growth of intestinal microbes on fiber, including fiber based prebiotics. In summary, we demonstrate that our Protein-SIP approach allows for the ultra-sensitive (0.01% to 10% label) detection of stable isotopes of elements found in proteins, using standard metaproteomics data.

## Introduction

Microbial communities drive chemical transformations from global element cycling to human nutrition. Unfortunately, the overwhelming complexity of these communities is often a barrier to unraveling their functionality. Use of isotopic or chemical labeling is a powerful solution to that problem. Even in the context of complex microbial communities, labeling enables assigning activities and functions to taxa, tracking metabolic pathways and resolving trophic relationships among species^[1–5]^. Current labeling approaches include use of click-chemistry (BONCAT)^[6]^, nanoscale secondary ion mass spectrometry (nanoSIMS)^[2]^, Raman microscopy ^[7]^, genomic sequencing of isotope labeled DNA/RNA (DNA/RNA-SIP)^[8]^, separated from unlabeled DNA/RNA with density gradient centrifugation, and protein-based stable isotope probing metaproteomics (Protein-SIP)^[9]^. Some of these approaches use labels with defined chemistry such as non-canonical amino acids in BONCAT^[6]^, which are directly assimilated into biomass. Others use more generic labels, such as substrate molecules labeled with heavy isotopes of carbon, nitrogen, oxygen and hydrogen^[2,7,10,11]^. When spatial organization of samples is important, approaches are available to image labeling outcomes^[12,13]^. When it is unknown in advance which species or pathway might be involved in a target process, labeling can be combined with untargeted metagenomics and metaproteomics analyses.

Recently, we developed an algorithm (Calis-p 1.0) to estimate natural isotope abundances (stable isotope fingerprints, SIF) of carbon isotopes of individual species within complex microbial communities using metaproteomics ^[14]^. In nature, ^13^C and ^12^C occur side by side at a ratio of approximately 0.011 ^13^C/^12^C. For microbial biomass, very subtle changes to this ratio, as little as 0.0001, already provide information about carbon assimilation pathways and carbon sources used. Our algorithm, which modeled mass spectra of individual peptides using Fast Fourier Transformations (FFTs), was able to detect these subtle changes. In the present paper we further develop this extremely sensitive approach to also work for stable isotope probing (SIP) experiments. This enables us to detect and quantify the assimilation of heavy isotopes by individual species in complex microbial communities using metaproteomics (Protein-SIP).

Protein-SIP differs from other metabolic labeling approaches in that the heavy isotopes from the substrate are incorporated into protein through *de novo* synthesis of amino acids from the substrates via biosynthetic pathways, rather than directly in the form of labeled amino acids. Such labeled amino acids are used, for example, in the “Stable Isotope Labeling by Amino Acids in Cell Culture” (SILAC) approach ^[15]^. The “random” incorporation of label into various amino acids and ultimately into peptides makes data analysis much more complicated in Protein-SIP, at least compared to the predictable exact mass shifts resulting from direct assimilation of labeled amino acids in SILAC.

Protein-SIP approaches have been successfully developed before, but these approaches have their challenges (for an overview see introduction of ^[10]^). Metaproteomics relies on high-resolution mass spectrometry to detect, identify and quantify peptides, which are then used for protein identification and quantification^[16]^. Using the same mass spectra already used for peptide identification to also quantify abundances of heavy isotopes in these peptides appears a straightforward add-on, as these spectra resolve the peptide isotopes and provide their intensities. However, unknown amounts of heavy isotopes shift peptide mass peaks by unknown numbers of mass-units, which makes the identification of peptides based on masses computationally challenging. The existing Sipros algorithm solved this problem with brute force by coupling the detection of labeled peptides with the initial peptide identification. Sipros predicts the most abundant peptide masses and isotopic distributions of b and y ions in an isotope atom% range of 0 - 100% in 1% increments^[17]^. This approach makes Protein-SIP experiments computationally so expensive that dedicated smaller protein sequence databases have to be constructed for determination of stable isotope content of peptides^[18]^ and even then the approach still requires a supercomputer to work. For example, one study using the Sipros approach had to invest around 500,000 CPU hours for a study with less than 10 labeled samples^[10]^. The MetaProSIP^[19]^ and SIPPER ^[20]^ algorithms overcame the problem by using spectra of unlabeled peptides as a starting point for computations. In case of MetaProSIP these unlabeled peptides can be derived from the SIP experiment itself if a portion of the original unlabeled proteins is still present, or, alternatively, from a control sample that was incubated without label. MetaProSIP then detects the labeled peptides corresponding to the unlabeled peptides and computes the relative isotope abundance and labeling ratio based on the comparison of the labeled and unlabeled form of peptides ^[19]^. Because MetaProSIP requires a labeled peptide’s spectrum to be shifted away from the mono-isotopic mass, it has been speculated that it requires relatively heavy labeling (e.g. >20.24 atom% for ^13^C and >73.1 atom% for ^15^N ^[21]^). In case of SIPPER the isotopic patterns for unlabeled peptides are generated in silico and subtracted from the experimental isotope patterns of peptides. Remaining peak intensities after subtraction are used for estimating isotope content. SIPPER is designed to detect small changes in isotopic profiles of complex mixtures after short exposure to 13C label, with proposed scoring schemes to reduce the rate of false discoveries.

While the identification challenges can be solved by clever algorithms, underneath these challenges hides a more fundamental problem. Figure 1 shows the expected mass spectra of three *E. coli* peptides after 1/8 generation of labeling with ^13^C-glucose. The figure illustrates the problem with these data: Assimilation of heavy isotopes into peptides leads to broadening of spectra. Thus, a peptide’s matter gets divided over ever more peaks, reducing sensitivity. Also, because many peptides get injected into the mass spectrometer simultaneously, especially for complex samples such as a microbial community, the probability of the peptide’s spectrum overlapping with another spectrum increases as it broadens, reducing data quality. Heavy peptides are especially sensitive to these issues. Counter-intuitively, for Protein-SIP, sensitivity is highest when using small amounts of label.

**Figure 1:**
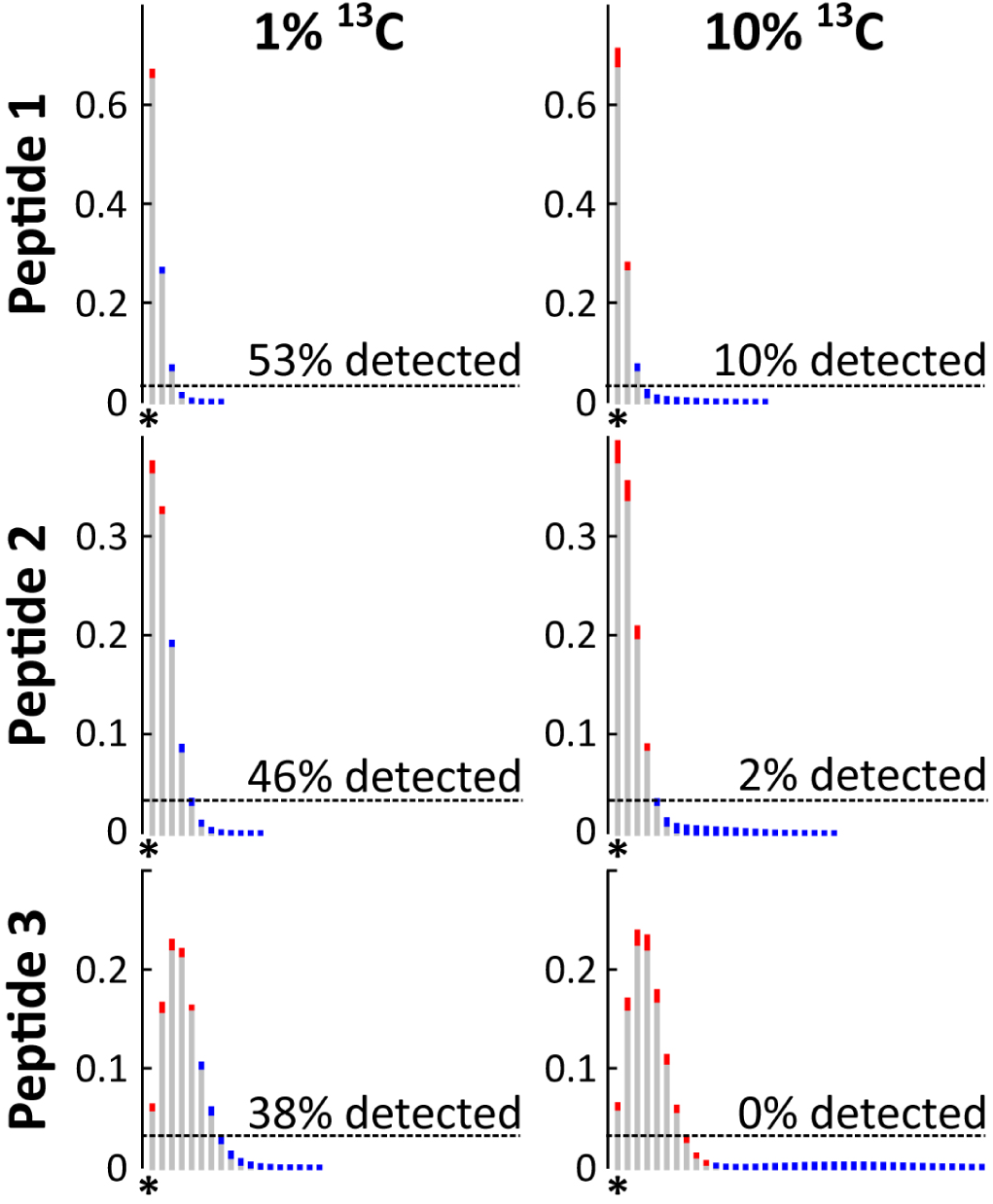
Modeled spectra of three *E. coli* peptides after ⅛ generations of growth on 1% (left) and 10% (right) ^13^C_1-6_ glucose (13C/12C 0.02 and 0.11 respectively). Assimilation of 13C into peptides leads to a shift of matter away from from the monoisotopic mass (shown as *). The resulting peak intensity changes are shown in red - for peaks with decreased intensity -, and blue - for peaks with increased intensity after labeling. Dashed lines show experimentally determined average detection limits for peaks (see methods). Peaks below the dashed line are not recorded by the mass spectrometer. Percentages above lines indicate how much of the actual change is detectable in practice. Peptide 1 - IGLETAR; peptide 2 - AFEMGWRPDMSGVK; peptide 3 - QIQEALQYANQAQVTKPQIQQTGEDITQDTLFLLGSEALESMIK.

Our previous algorithm was developed to estimate slight differences in isotopic content based on peptide mass spectra, to determine natural carbon isotope abundances^[14]^. It made use of the fact that in nature, heavy isotopes are distributed randomly, yielding spectra that are perfect Poisson distributions. This enabled us to reduce the noisiness of the data by identification and rejection of imperfect spectra. Spectra in Protein-SIP experiments do not have such conveniently predictable properties. With labeled samples, the shape of spectra cannot be predicted using FFT, because these spectra become mixtures of spectra associated with labeled and unlabeled peptides. Both the proportion of heavy isotopes in the labeled peptides and the extent of labeling - the relative abundances of labeled versus unlabeled populations of peptides - are unknown in advance. For analysis of these data we therefore developed rigorous noise filtering and estimated isotopic content based on neutron abundance, requiring no assumptions about a spectrum’s shape.

We present new algorithms and software for sensitive and quantitative estimation of isotopic content of individual species in stable isotope probing experiments with complex microbial communities. The new algorithms have been integrated into the Calis-p software together with the SIF algorithms, and the software was completely re-written to enable Protein-SIP (new version is Calis-p 2.1). The software decouples peptide identification from label detection and is thus compatible with most standard peptide identification pipelines.

Computation of label content is very fast, a high-end desktop computer only needs one minute for processing ∼1 Gb of data, corresponding to ∼10,000 MS1 spectra or ∼40 min of Orbitrap runtime. Using pure cultures of bacteria and mock communities, we show that Protein-SIP with Calis-p yields best results when substrates are partially labeled. For example, for carbon the fraction of heavy atoms should make up <10% of the total. For abundant organisms, assimilation of label (such as ^13^C) into protein can be quantified within minutes after adding the label, within 1/16 of a generation. Even for rare organisms making up ∼1% of a community, a single generation of labeling is sufficient for robust detection of label assimilation. We believe these advances will be helpful to microbiome researchers and microbial ecologists seeking to assign functions and activities to taxa, to track metabolic pathways and for resolving trophic relationships among species.

## Results

Previously we presented algorithms and software for estimating natural isotope fingerprints from peptide mass spectra^[14]^. Our previous algorithm made use of the stochastic distribution of isotopes in nature and mass spectra that can be modeled by Fast Fourier Transformations. Quality control is intrinsic to that approach, as poor quality spectra cannot be modeled with FFT and can be rejected. Examples of low quality spectra are spectra that overlap with other spectra or low intensity spectra that are affected by noise. Feeding microbes labeled substrates for Protein-SIP experiments leads to peptide mass spectra with irregular shapes that cannot be modelled with FFT, as explained in the introduction. Isotopic composition of such spectra can still be inferred, by adding up the mass intensities of all peaks in the spectrum according to Equation 1 in the Methods (implemented as “neutron abundance” model in Calis-p). Unfortunately, that approach does not enable rejection of low quality spectra. Therefore, we implemented a simple noise filter based on unsupervised Markov clustering of all spectra associated with a single peptide (see Materials and Methods for details). The assumption underlying this approach is that most spectra are relatively unaffected by noise and will form the largest cluster. Spectra outside the largest cluster should be rejected for being of lower quality.

The performance of this filter was benchmarked using previous natural-isotope abundance data of pure cultures and mock communities of microbes (Suppl. Results & Discussion, Fig. S1, Tables S1 & S2). The FFT estimates of ^13^C/^12^C ratios for filtered spectra was as good or better than reported previously without filtering^[14]^. Even better, after filtering the estimates of ^13^C/^12^C ratios according to Equation 1 (see Materials and Methods) were now almost as good as for FFT. The average difference between the actual and estimated median δ^13^C values for the fifteen most abundant organisms of a mock community was 2‰ for FFT and 4‰ for the neutron abundance model. Implementation of the filter also dramatically reduced computation times because fewer FFT operations were needed.

To test the performance of Equation 1 for ^13^C labeled peptides, we labeled cells of two model organisms, *Escherichia coli* K12 and *Bacillus subtilis* ATCC 6051, to saturation, with ^13^C glucose. Three replicate cultures were grown overnight with fully-labeled glucose (^13^C_1-6_) and single-labeled glucose (^13^C_2_), with the percentage spiked-in ^13^C increasing from 0 and 10% of total glucose in seven steps. The glucose that was used as unlabeled glucose had the natural ^13^C content of around 1.1%. Protein was extracted, peptides were prepared, subjected to LC-MS/MS, and identified with SEQUEST HT in Proteome Discoverer and results analyzed in Calis-p (see Materials and Methods).

Surprisingly, adding as little as 1% label could already severely compromise the identification of peptides by search algorithms such as SEQUEST. For example, for *B. subtilis* average peptide-spectrum matches (PSMs) dropped by almost 90% at 10% added label. To mitigate these losses, we tested five potential improvements to search strategies (see Materials and Methods and Supplementary Results & Discussion for details). These ranged from computationally costly, open mass window searches to more confined, faster approaches. All strategies improved identification outcomes (Fig. 2 & S2). However, for the “open search” and “dynamic modifications” strategy, computation times became impractical. For example, basic peptide identification on a high performance desktop computer with the SEQUEST algorithm for single 140 min LC-MS/MS file from a microbial community took 17 min to search with our standard search parameters, while it took 72 min with a search strategy that included modifications of the peptide termini, and 1,947 min for a search strategy with a open (20 Da) mass window. We selected the Modifications of Termini (N=low, C=high) strategy as a practical compromise between peptides identified and extra computation time needed. This strategy adds six custom “post-translational” modifications to the protein identification search. These modifications generated more PSMs by enabling addition of one to three neutron masses at the N-terminus of a peptide and four to six neutron masses to its C-terminus. We observed a strong difference in how label amount impacted the number of PSMs between B. subtilis and E. coli. For B. subtilis a small amount of added label strongly increased the number of PSMs, which then sharply dropped at 1% label. We currently have no good explanation for this phenomenon.

**Figure 2:**
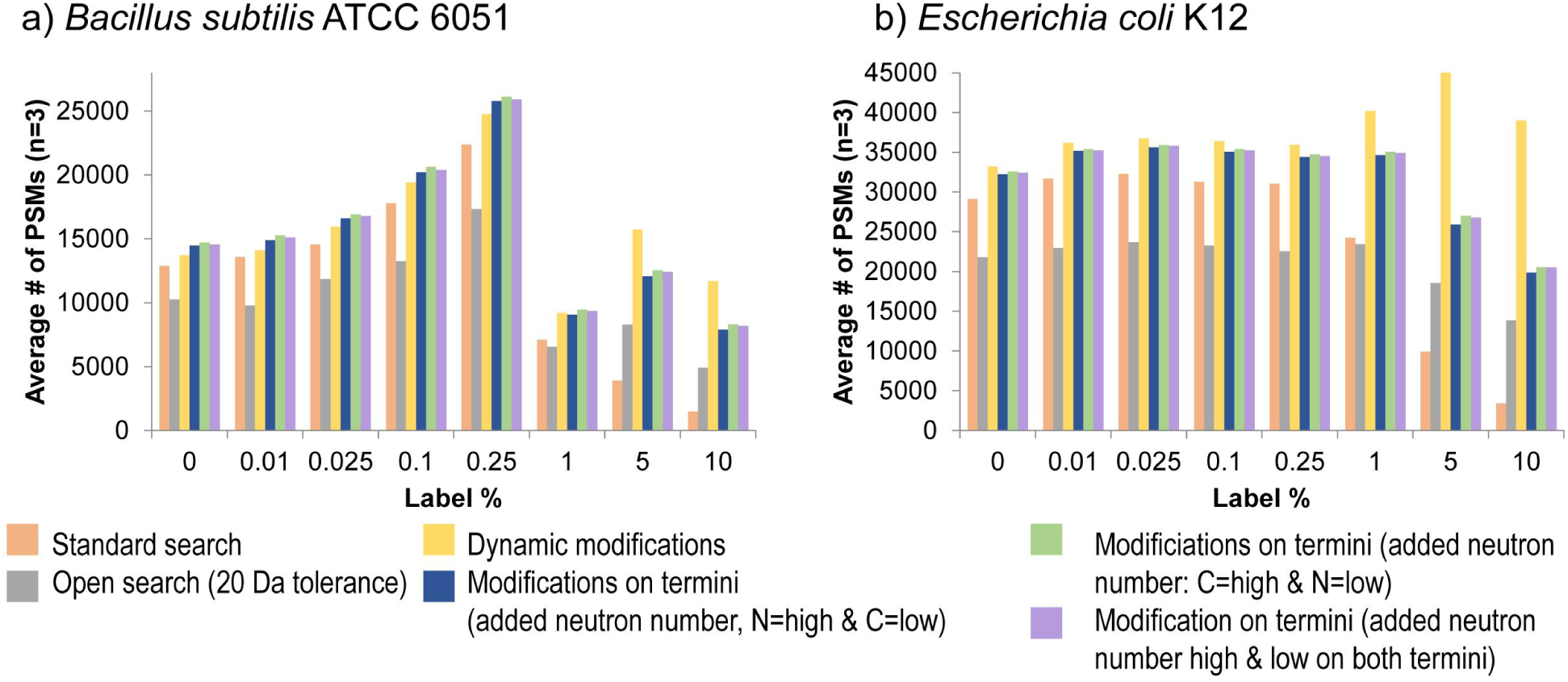
A small modification of the peptide identification approach drastically increases the number of peptides with 1-10% label that can be identified. Number of peptide spectral matches (PSMs) identified at different ^13^C label percentages using six different peptide identification strategies. Cultures of a) *B. subtilis* and b) *E. coli* were grown in Bacillus minimal medium or M9 minimal medium (*E. coli*) in which a percentage of the glucose was replaced with ^13^C_1-6_ glucose for >10 generations to achieve close to complete labeling. Three biological replicates were run for each label percentage. Peptides were identified using the SEQUEST HT Node in Proteome Discoverer (version 2.2) with six different strategies to account for the mass shifts caused by addition of heavy atoms. Standard search: no dynamic modifications to account for addition of label; Open search: the precursor mass tolerance was set to 20 Da allowing for the potential addition of 20 neutrons (e.g. ^13^C atoms) in a peptide; Dynamic modifications: allowing for up to three dynamic modifications each of two custom peptide modifications adding a 1 neutron mass shift and a 2 neutron mass shift (up to 9 neutrons in total per peptide); Modifications on termini: six dynamic modifications were set up that were restricted to either the C or the N-terminus of the peptide. The modifications account for mass shifts of 1 to 6 neutrons and depending on the search strategy the low mass shifts (1, 2 and 3 neutrons) were set up as modifications on the C or the N-Terminus or low and high mass shift modifications were distributed between both termini. Each modification can only be added to a terminus once. This strategy allows for a total of 21 neutron additions to a peptide.

After assigning peptides to spectra with the improved peptide identification strategy, low quality spectra were rejected using the filter described above. For the remaining spectra, the number of neutron masses added as custom modifications during the identification step already provided a qualitative, or at best semi-quantitative, measure for label incorporation (Fig. 3, Suppl. Table S3). However, inference of the ^13^C/^12^C ratios by Equation 1 was much more precise, even for minimally (0.01%) labeled cells providing a limit of detection <0.01% label in most cases (see supplementary text). Precision and especially label recovery were both higher when using glucose labeled at only a single position rather than with fully labeled glucose. For the latter, the recovery was only 75-79%, meaning that the ^13^C/^12^C ratio was 21-25% lower than expected. Potentially, this was caused by broadening of spectra with fully labeled glucose (Fig. 1). As explained in the introduction, broader spectra reduce sensitivity. Interestingly, the breadth of spectra could be used to infer to what degree ^13^C carbon was assimilated in clumps of multiple atoms (pie charts in Fig. 3). This approach, which only works when all atoms in a substrate are labeled and when cells are labeled to saturation, could be used to infer the number of carbon atoms in substrates that a given species is assimilating. In other words, Protein-SIP can provide hints on whether a species is autotrophic or heterotrophic. We carried our similar labeling to saturation experiments with 15N (E. coli labeled to saturation with 2.5% 15N ammonium) and obtained similar results (data not shown, but available via PRIDE see data accessibility statement).

**Figure 3:**
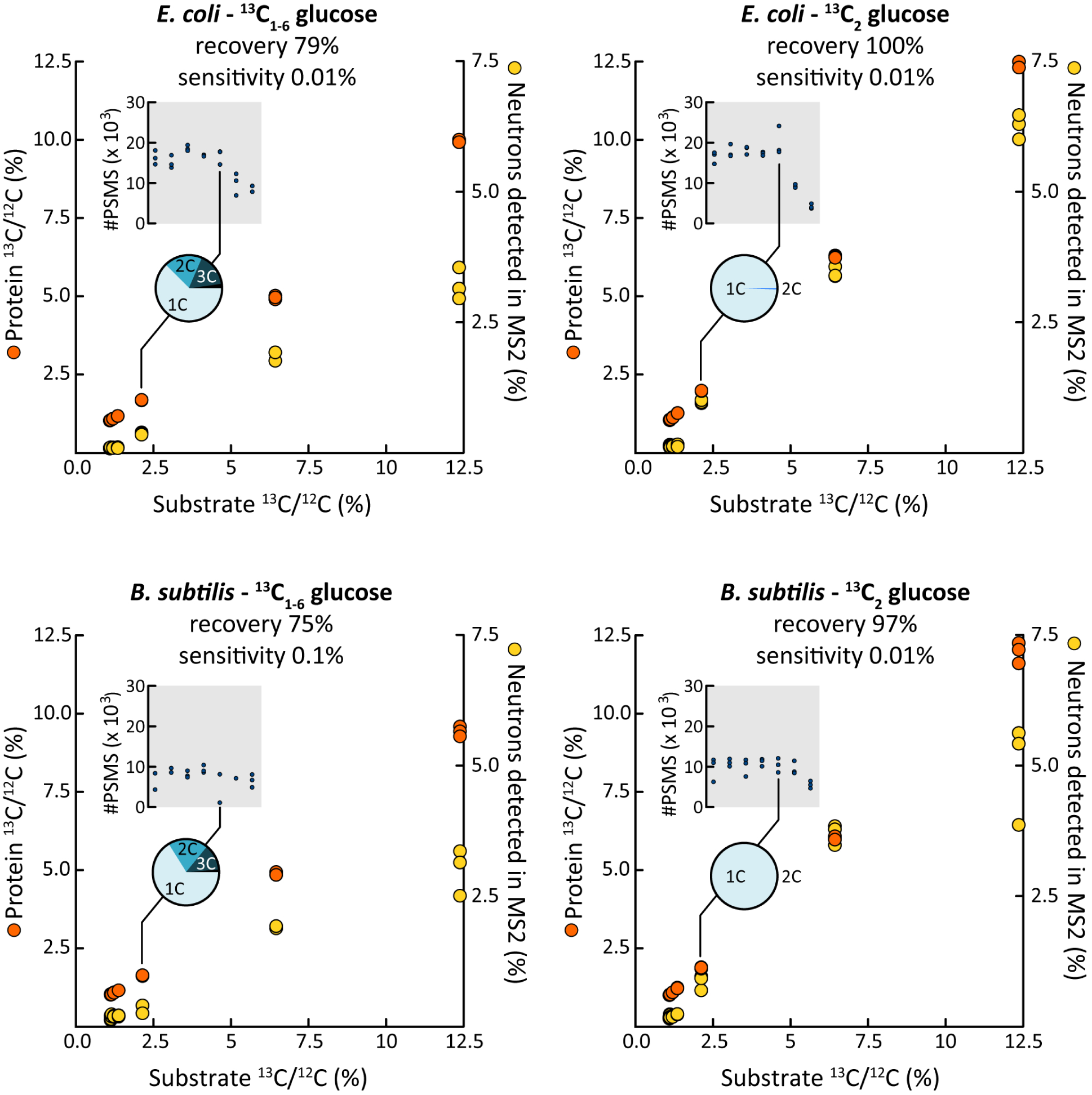
The number of labeled atoms per substrate molecule impacts the ability to quantify label incorporation accurately. Labeling, to saturation, of *E. coli* and *B. subtilis* with single-labeled (^13^C_2_) and fully labeled (^13^C_1-6_) glucose. The ^13^C/^12^C ratio in the substrate was varied. Note that unlabeled glucose (0% added ^13^C glucose) has a natural ^13^C content of around 1.1%. Each orange circle is the median ^13^C/^12^C ratio of all peptides measured in one replicate incubation (on average 2758 peptides per replicate). Determined 13C/12C ratios increased linearly with substate ^13^C/^12^C ratios (R^2^ >0.999). Almost 100% of the substrate ^13^C was recovered in protein for ^13^C_2_ glucose labeled cells. Recovery was lower for ^13^C_1-6_ glucose. The proportion of neutron masses detected via the improved peptide identification strategy using N- and C-terminal modifications (yellow circles) increased with substrate ^13^C/^12^C ratios, but at low linearity and sensitivity. The number of Calis-p filtered Peptide Spectrum Matches (PSM) decreased for ^13^C/^12^C ratios above 2.5% (insets) as expected based on Figures 2 & S2. Assimilation of carbon into amino acids in clumps of multiple ^13^C atoms was detectable in peptide spectra of cultures fed with ^13^C_1-6_ glucose as shown in pie charts for experiments fed with ^13^C/^12^C 1% above natural background. The detailed data for this figure can be found in Suppl. Table S3.

The data of Figure 3 are not yet a meaningful representation of what an actual Protein-SIP experiment would look like. In practice, we would always avoid labeling a microbial community to saturation, because all community members would end up being labeled equally, providing no new information on elemental fluxes and substrate uptake in the community. To mimic an actual Protein-SIP experiment, we mixed labeled and unlabeled cells of *E. coli* at different ratios, leading to compound spectra as shown in Figure 1.

The results indicated that estimation of ^13^C/^12^C ratios with this type of compound spectra was more challenging (Fig. S3, Suppl. Table S4). Also, the difference between single labeled and fully labeled glucose was more pronounced, with the former yielding much better sensitivity and label recovery than the latter. For example, at 1% label recovery was 92% for single labeled glucose, while it was 80% for glucose with 6 labeled carbons.We also compared the performance of two center statistics for ^13^C/^12^C ratios, the intensity-weighted mean and the median. The intensity-weighted mean displayed higher sensitivity and precision than the median in these experiments (for contrasting results for community samples see below). However, both with 1% and 10% single labeled glucose, even the median ^13^C/^12^C ratios accurately quantified label assimilation within 1/16 of a generation (simulated by mixing of labeled and unlabeled cells), corresponding to as little as 1-2 min of growth for *E. coli*.

Next, we investigated whether our approach was capable of detecting label assimilation in the context of a microbial community. For this, we used a previously described mock community, comprising >30 microbes, including gr+ and gr-bacteria, an archaeum, a eukaryote (algae) and several phages^[22]^. This community also included *E. coli* K12, at ∼6% abundance. Here, we mixed cells of *E. coli* labeled with 1%, 5% and 10% ^13^C_1-6_-glucose into the unlabeled mock community at a ratio corresponding to one generation of growth for *E. coli*. Quantification of ^13^C assimilation was straightforward and linear (R^2^ 0.99, Fig. 4A, Suppl. Table S5). This was perhaps not surprising because a relatively large amount of label was used and the relative abundance of *E. coli* in the mock community was high, i.e. ∼12% after addition of the labeled cells.

**Figure 4.**
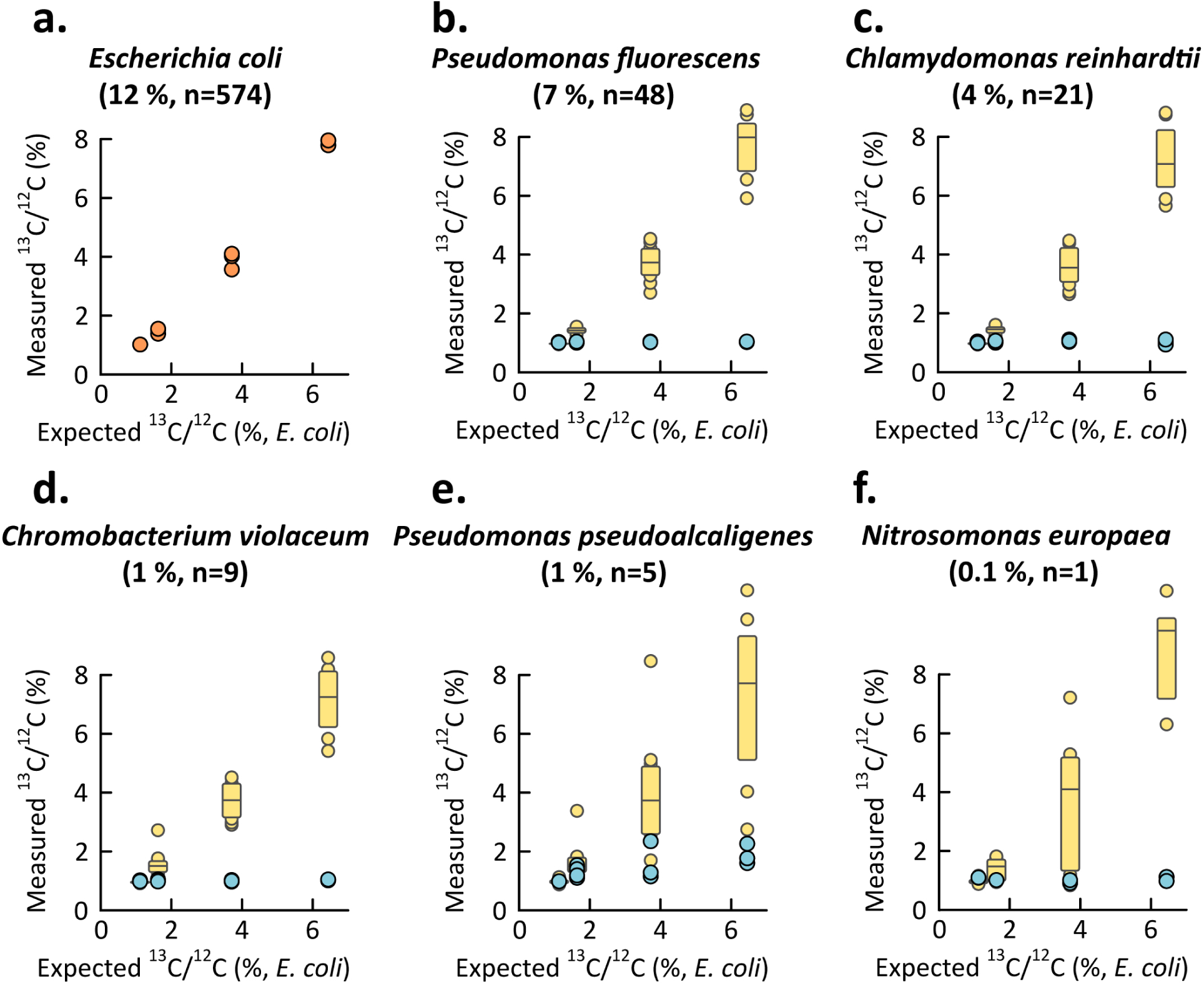
Detection of ^13^C assimilation by *E. coli* within a mock community of 32 microorganisms developed by ^[22]^. In each experiment, half of the *E. coli* cells were labeled using ^13^C_1-6_-Glucose, corresponding to one generation of labeling, with glucose containing 0, 1, 5, and 10% ^13^C on top of natural abundance ^13^C (three replicate samples were generated for each labeling percentage and measured separately). Label incorporation by *E. coli* (**orange circles in a**), but not by other organisms (**blue circles shown for five organisms in b-e**), was clearly detectable and reproducible. Yellow box plots show the measured ^13^C content of sets of *E*.*coli* peptides, obtained by downsampling of the results in **a**, mimicking the spectral intensities of the peptides collected for each unlabeled organism in panels **b-e** i.e. only *E. coli* peptides that corresponded in intensity to peptides of the analyzed organism were used. The percentage in parentheses indicates the relative abundance of the organism in the mock community based on its proteinaceous biomass and the “n=“ indicates the average number of peptides passing the filters in Calis-p for SIP value calculation for the organism in each experiment, which also corresponds to the number of *E. coli* peptides used in downsampling. These results show label incorporation can be estimated, even for relatively rare species. Suppl. table S5 shows results for each species.

Figure 4 also shows to what extent the addition of the labeled cells of *E. coli* led to the incorrect inference of label assimilation by five other members of the mock community. The relative abundance of these unlabeled organisms was between 0.1% and 7%. The determined ^13^C/^12^C ratios for the >20 other members of the mock community are reported in supplementary table S5. We found that the choice of center statistic used has a major impact on the false positive detection of label incorporation. When using the median, the overall (i.e. all unlabeled species in all replicates) False Positive Rate (FPR) of label detection for populations with nine or more peptides (after filtering) was 3.4% and for populations with eight or fewer peptides it was 45%. In contrast, when using the weighted mean the FPR was 51% for populations with nine or more peptides and 50% for populations with eight or less peptides. In our dataset, the nine peptide threshold corresponded to ∼1% relative abundance of strains/species within the mock community. We investigated the massive differences in FPRs between the two center statistics by manually checking spectra causing false positives and found that low-intensity peptide spectra associated with less abundant populations were often affected by the overlap with broadened spectra of a labeled, more-abundant population. Therefore, we concluded that for label detection in microbial communities the median should be used (see detailed discussion in supplementary methods). Figure 4 shows examples of false-positive inferences for *Pseudomonas pseudoalkaligenes*.

To investigate whether label assimilation can be correctly inferred for less abundant populations, we downsampled (bootstrapped, up to ten times) the set of >6,000 peptides collected for *E. coli*, using the peptides of each other organism as templates. In the resulting datasets, each *E. coli* peptide was matched to a peptide of the other organism with a similar intensity. Based on inferences for these bootstrapped datasets shown in Figure 4, label assimilation could be robustly estimated, at least for populations associated with nine or more peptides, corresponding to ∼1% abundance. This number of peptides is much smaller than the ∼30 peptides needed for estimation of natural carbon isotope content in a species using Protein-SIF (Suppl. Results & Discussion, Fig. S1).

Next, we analyzed how well we could detect incorporation of label into individual proteins based on how many peptides passed the Calis-p quality filters for a protein. For this we analyzed the Calis-p reported ^13^C/^12^C ratios for proteins from the mock communities with 5% labeled *E. coli* spiked-in and without spiked in *E. coli*. ^13^C/^12^C ratios in *E. coli* proteins from the 5% spike-in samples were on average much higher than the ratios for proteins from the unlabeled mock communities and the unlabeled mock community members in the 5% spike-in samples (Fig. 5a). Even for proteins for which only 1 peptide passed the Calis-p quality filters, this pattern was observed. This indicated that label incorporation into individual proteins can be detected with as few as 1 peptide. For some proteins from unlabeled organisms, for which 3 or less peptides passed the Calis-p quality filter, ^13^C/^12^C ratios that were above the expected value of 0.011 (Fig. 5a), indicating that, as expected, the accuracy of the ratio estimates increases with more peptides available per protein.

**Figure 5:**
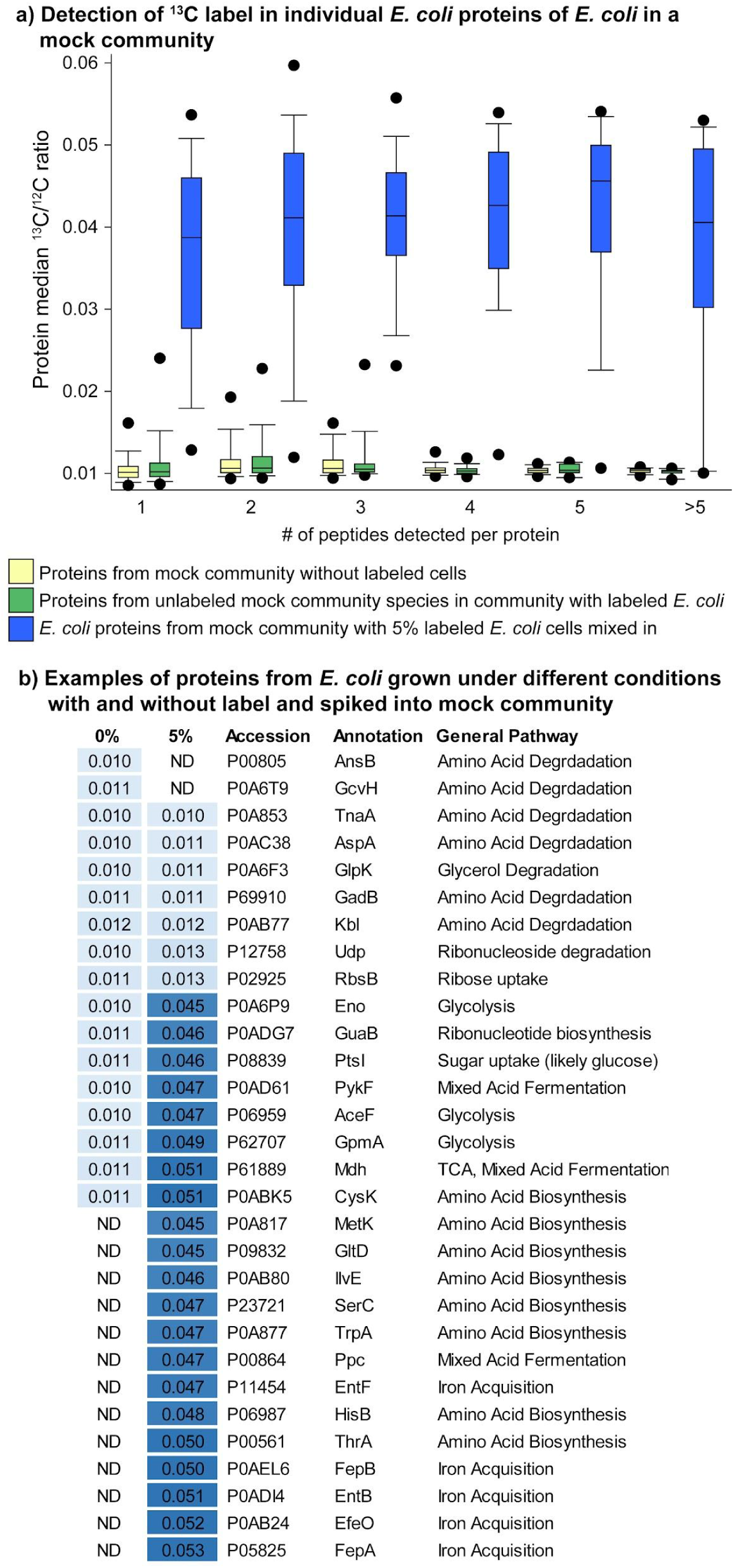
Measurement of ^13^C label content in individual proteins. Analysis of a subset of the data shown in Fig. 4. *E. coli* grown in standard LB medium without label (0% added label) was part of a mock community consisting of 32 microorganisms ^[22]^. To this mock community *E. coli* grown in minimal M9 medium with glucose (5% of total glucose as ^13^C_1-6_-Glucose) in air tight bottles under oxygen limiting conditions was added in a 1:1 ratio to the unlabeled LB grown *E. coli* cells in the mock community. **a)** Detection of increased ^13^C/^12^C ratios in individual proteins as a function of the total number of different peptides detected for each protein. Proteins from all species in the unlabeled mock community are compared to the proteins of all unlabeled species in the mock community that contained the 5% labeled *E. coli* cells, as well as to the proteins from *E. coli* in the mock community that contained the labeled *E. coli* cells. The boxes indicate the 25th and 75th percentile, the line the median, the whiskers the 10th and 90th percentile, and the dots the 5th and 95th percentile. **b)** Examples of *E. coli* proteins that showed no or high label incorporation in 5% ^13^C glucose grown *E. coli* in the mock community. Unchanged ^13^C/^12^C ratios shown in the table between treatments indicate that proteins were not produced in cells that were grown in M9 medium with labeled glucose, but were present in cells grown in LB. Proteins with high ratio were mostly or exclusively produced by cells grown in M9. ^13^C/^12^C ratios in the table are averages of three replicate samples. Only proteins that were detected in at least two replicates in one of the conditions are shown. The full table is Suppl. Table S6.

To test if meaningful results can be obtained from populations in mixed communities that shift their metabolism and physiology we analyzed ^13^C/^12^C ratios in *E. coli* proteins from the 5% spike-in samples in more detail. For this analysis it is important to know that the unlabeled *E. coli* cells that were part of the mock community were grown at well oxygenated conditions in a complex medium containing organic nutrients such as amino acids and vitamins (LB broth), while the *E. coli* grown in the presence of ^13^C_1-6_-glucose (5% of total glucose) were grown under oxygen limited conditions in a minimal medium (M9 broth) that contained glucose as the only carbon source and nitrogen only in inorganic form. This mixing of unlabeled LB grown and labeled M9 grown *E. coli* led to the following expectations: 1) proteins that are produced exclusively or almost exclusively by cells growing on LB would show no label incorporation in the 5% label mock communities and thus have ^13^C/^12^C ratios close to the expected value of 0.011; and 2) conversely proteins that are produced exclusively or almost exclusively by cells growing on M9 would show high label incorporation in the 5% label mock communities (^13^C/^12^C ratios of >0.04). We only looked at proteins that were detected in at least two replicates of one condition (Suppl. Table S6, Fig. 5b). Proteins for the degradation of amino acids and other carbon sources were unlabeled, indicating that they were only produced by *E. coli* in complex medium, but not when growing on M9. Proteins for amino acid biosynthesis pathways, glycolysis, mixed acid fermentation and iron acquisition were heavily labeled, indicating that lack of amino acid sources in the medium led to expression of biosynthesis pathways, oxygen limitation led to induction of fermentation pathways and that potentially cells growing in M9 were iron limited.

### Comparison with existing Protein-SIP approaches

To compare the performance of Calis-p with existing Protein-SIP approaches four of the above described datasets with labeled E. coli spiked into a mock community were processed by expert operators of the SIPPER and MetaProSIP workflows. We would like to highlight here that these two approaches were developed for higher label amounts (MetaProSIP) and low level labeling after shot label exposure (SIPPER) and they may well outperform Calis-p under specific conditions. However, we focused our comparison on the low label amounts for which the ultra-sensitive Protein-SIP within Calis-p was developed and a comparison at high label amounts (>10% label) was outside the scope of this study.

The three approaches differed strongly in the number of peptides for which label content was quantified (Fig. 6b). Part of this was due to the fact that for each of the three approaches different peptide identification algorithms were used (Fig. 6b) leading to differences in the number of peptides that served as input for label quantification. The MSGF+ search engine used by SIPPER yielded the lowest number of identified peptides. While MetaProSIP quantified label content for the highest proportion of input peptides in all cases (>80%), Calis-p provided isotope contents for the lowest proportion of input peptides (25-31%). This large difference for identified and isotopically quantified peptides for Calis-p is caused by the multiple quality filtering steps in Calis-p that remove large numbers of peptides and spectra from consideration for various quality related issues (e.g. flagged for No valid PSM’ ‘too few spectra’ ‘No majority vote while clustering’). With the higher number of peptides for which 13C content was quantified by MetaProSIP, MetaProSIP provided sufficient peptide numbers (>9 for a species) to quantify label content for the highest number of species (Fig. 6, Fig. S4).

**Figure 6:**
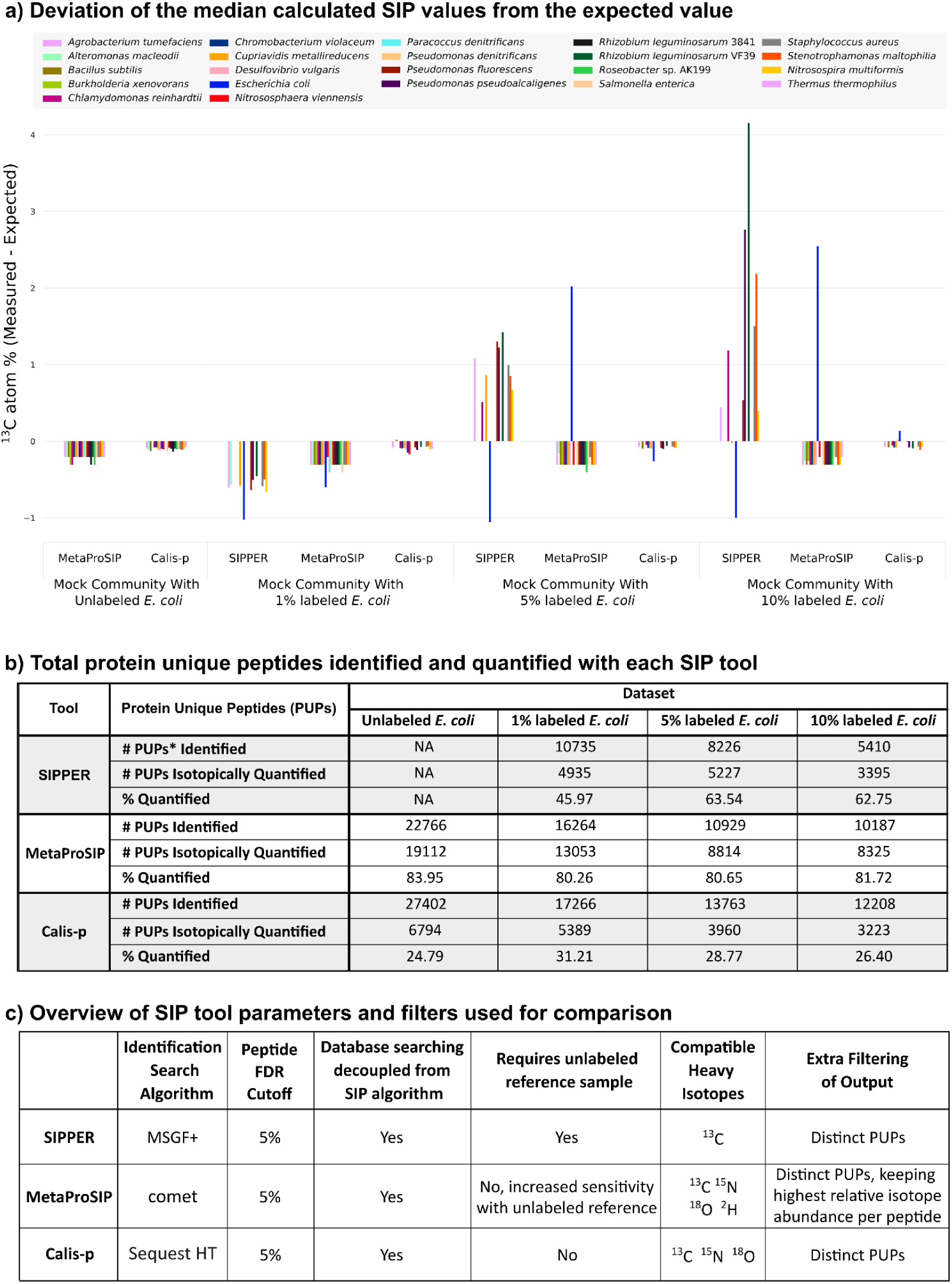
Comparison of the output from the three Protein-SIP approaches: SIPPER, MetaProSIP, and Calis-p. Four datasets were processed by expert operators for each approach using optimal parameters for each approach. The outputs from each tool were filtered for comparability by retaining only distinct protein unique peptides (PUPs), defined as peptides unique to a protein sequence and with a unique combination of sequence, charge state, and m/z. a) Median 13C values were determined for organisms with 9 or more peptides. The expected 13C atom % value for each experimental condition was subtracted from each experimental SIP value and the deviation of the experimental value from the expected value is displayed. b) Table showing the total number of protein unique peptides identified and used as the input for each approach and the total for which isotope values were quantified. c) Summary of the parameters used for each tool/approach and additional post-processing steps as recommended by each expert operator. Each tool output was filtered for distinct protein unique peptides i.e. isotope values were only used if the peptide could be uniquely assigned to a single species. MetaProSIP required an additional post-processing step for selecting the highest relative isotope abundance (RIA) value in cases where the tool reported multiple RIA values. Detailed data for this figure is shown in Figure S4 and tables S7 to S10.

The three approaches also differed in the number of false positive detection of above natural abundance 13C in peptides (Fig. S4, Tables S7 to S10) and species (Figs. 6a & S4). Calis-p overestimated 13C content only in a small number of peptides and in none of the species (i.e. median 13C content was at or below expected value). MetaProSIP also did not overestimate label content in any of the species, instead it had the tendency to underestimate isotope content of unlabeled species.

SIPPER had a high rate of false positive detections, while for the 1% labeled sample all 13C content estimates (including the estimate for the labeled E. coli) were strongly underestimated. 13C content of unlabeled species for which enough peptides were quantified (9 peptides) were strongly overestimated. This false positive label detection is likely due to the underlying principle of SIPPER, which is to identify labeled peptides, while not trying to classify unlabeled peptides. This means that truly unlabeled peptides are not available for calculation of label content of species and a small number of false positive label detection in peptides can lead to miss estimation of label content in species and proteins.

Finally, the three approaches also differed in the accuracy of 13C content estimates for the labeled E. coli. For the 1% labeled E. coli Calis-p was the only approach to detect the label in E. coli with the median 13C content being close to the expected value (Figs. 6 & S4) indicating a higher sensitivity of Calis-p at low label contents. For the 5% and 10% labeled E. coli all three approaches detected label in E. coli with the median 13C values being closest to the expectation for Calis-p in both cases indicating a higher accuracy of the Calis-p estimate.

In summary, while MetaProSIP outperformed Calis-p in terms of quantity of peptide isotope quantification, Calis-p was more sensitive and accurate. The sensitivity of MetaProSIP was, however, much better (detection down to at least 5% 13C) than recently suggested by Starke^[21]^. Both MetaProSIP and Calis-p showed high specificity; the lower specificity shown by SIPPER was likely caused by SIPPER’s primary focus on the detection of labeled peptides. Finally, while all three tools are independent of the search engine (in contrast to Sipros), Calis-p has been implemented to work with the open search engine output format mzIdentML, which will minimize compatibility issues between search engine output and Calis-p.

### Case Study: Differential heavy water incorporation reveals activity changes for intestinal microbiota species in response to dietary changes

To demonstrate the power of the Calis-p approach and to test our approach for additional elements we analyzed data from a complex microbial community grown with two types of heavy water^[11]^. Both 2H and 18O water can function as markers of translational activity[10,23] and thus allow to detect activity changes in members of microbial communities in response to changes in environmental conditions or use of complex substrates without relying on direct labeling with a specific substrate. The community consisted of 63 species isolated from human fecal material and was grown in bioreactors with either a high fiber medium or a high protein medium to simulate different dietary conditions encountered by the intestinal microbiota. For each diet treatment three replicate cultures were grown for 12 hours with unlabeled water and water in which 25% were either replaced with ^2^H_2_O or H_2_^18^O. We obtained thousands of peptides passing Calis-p quality filtering for each sample (mean 4602 peptides/sample, SD 1140, Fig. S5) and we were able to quantify heavy water incorporation (>9 peptides passing Calis-p filters for species) for 21 to 30 species per sample (Mean = 27, Median = 28). We found that overall incorporation of ^18^O was much higher than incorporation of ^2^H (Fig. S4). The low measured incorporation of ^2^H can potentially be attributed to variation in retention times of isotopically different forms of a deuterated peptide in reversed-phase chromatography^[24,25]^. Such retention time variation can generate distinct isotope patterns for the same peptide at different retention times, which would lead to failure to cluster by the Markov clustering in Calis-p during spectrum filtering. Since the amount of 2H used in this experiment was relatively high we do expect relevant retention time shifts of deuterated peptides. Additional factors that might explain low measured incorporation of ^2^H are the known strong fractionation of hydrogen isotopes in organisms^[26]^, the fact that many hydrogen atoms on peptides can freely exchange with water leading to loss of label during sample preparation^[27]^, the dilution of 2H in stable C-H bonds in de novo synthesized amino acids by hydrogens derived from organic growth substrates^[23,28]^, and the known toxicity of deuterium to many organisms slowing down their growth rates and thus reducing the rate of incorporation, which however usually occurs at higher concentrations (>50%) of deuterium than used in this experiment^[23,29]^. Finally, incorporation of ^18^O over ^2^H may be favored because for incorporation of ^18^O into non-exchangeable positions amino acid *de novo* synthesis is not required. ^18^O can be incorporated into the carboxyl group of amino acids during proteolytic cleavage of substrate proteins^[30]^ and remain in the peptide bond upon formation of new peptide bonds. While ^18^O in peptide bonds is stable and does not freely exchange with water, ^2^H in many positions on the peptide readily exchanges with H from water^[27]^. For hydrogen to be in positions with low exchangeability, amino acid *de novo* synthesis is required, because the necessary carbon-hydrogen bonds are only generated then^[27]^.

On the whole community level, label incorporation was significantly higher in communities grown with high protein as compared to high fiber (Fig. S5). On the level of single species we observed that responses to a change in “diet” were species-specific with some species such as *Akkermansia muciniphila, Bacteroides ovatus*, and *Clostridium bolteae* incorporating significantly more label under high protein conditions, while other species, such as *Alistipes onderdonkii, Clostridium lavalense*, and *Flavonifractor plautii*, showed no change or non-significant trends towards higher incorporation under high fiber conditions (Fig. 7). While incorporation trends for species between ^2^H and ^18^O were mostly consistent with each other, much fewer comparisons tested significant for ^2^H (4 for ^2^H versus 9 for ^18^O) likely due to the overall low measured incorporation of ^2^H and resulting low sensitivity. Our results indicate that availability of higher amounts of protein increases the translational activity of many intestinal microbiota species, which is in line with previous studies showing nitrogen, and by extension protein, is the limiting nutrient for the intestinal microbiota^[31,32]^. This indicates that mixed community bioreactors can be a useful analog to the intestinal tract for studying specific ecological factors (such as nitrogen limitation) driving community function. Surprisingly, although typically described as fiber degrading specialists^[33–35]^, we saw a significant increase of activity in several *Bacteroides* species in the high-protein medium relative to the high-fiber medium. This shows that it is critical to assess nitrogen/protein supply when analyzing fiber dependent growth of intestinal microbes. Furthermore, it suggests that nitrogen/protein supply is critical to consider when developing fiber based prebiotics to manipulate intestinal microbiota species^[36,37]^, which to our knowledge has not been considered so far. In summary, our results show that the use of heavy water for Protein-SIP allowed us to detect changes in the activity of microbiota members in response to changes in complex substrates.

**Figure 7:**
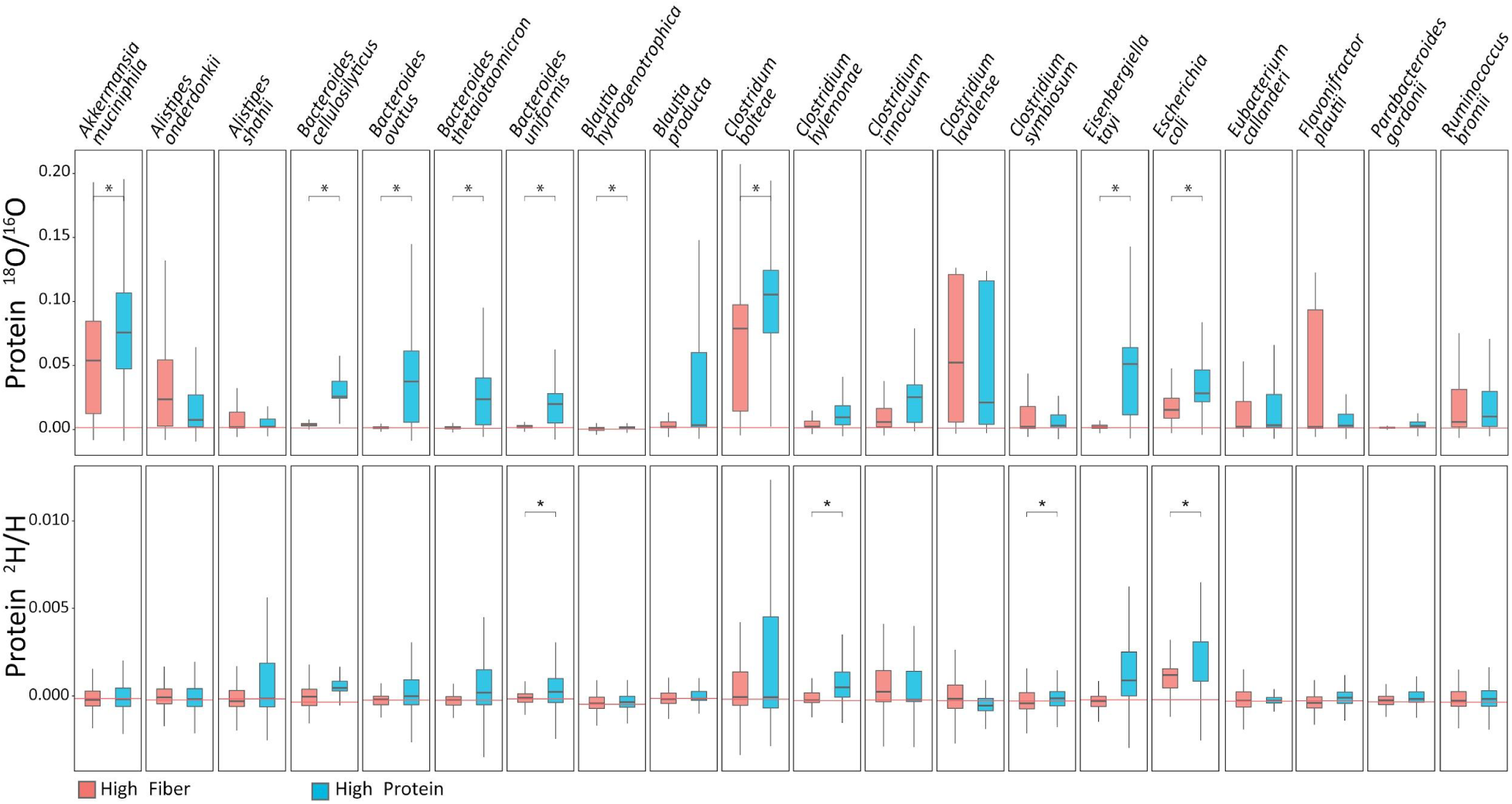
Strong differences in heavy water incorporation in intestinal microbiota species in response to diet. 63 species isolated from human intestinal microbiota were grown together in triplicates in either a high fiber or high protein medium in the presence of unlabeled water or water with either 25% ^2^H or ^18^O^[11]^. Calis-p based stable isotope ratios are shown for the 20 species for which at least 9 peptides passed Calis-p filtering conditions in all replicates. Each box shows the data for all peptides of the triplicate cultures combined (27 to 2225 peptides per box). The red lines indicate the average median for each species in the control samples with unlabeled water. Statistically significant differences are indicated with ‘*’ based on Student’s t-test on the means of replicates at p<0.05.

### BOX: Summary of Practical Workflow Considerations

The Protein-SIP workflow consists of several steps including (1) incubation of a microbial community with a isotopically labeled substrate, (2) metaproteomic sample preparation and LC-MS/MS data acquisition, (3) peptide identification, (4) data conversion and input to Calis-p, (5) isotope pattern extraction and computation of isotope content in Calis-p, and (6) analysis and interpretation of data provided by Calis-p (Fig. 8). The provision of isotopically labeled substrates in experiments can take many forms, such as addition of substrate to incubations of enrichment cultures/bioreactors^[11]^, addition to animal feed^[38,39]^, CO_2_ in plant incubation chambers^[40]^ or as ^15^N in plant fertilizer, and *in situ* incubations^[10]^. For the Protein-SIP approach presented here, substrate should be supplied with 1-10% of the total substrate containing the heavy isotope (label). Please note that this range refers to ^13^C, for other elements, such as N, which make up a smaller portion of atoms in a peptide, a higher amount of label can be used, as the associated peptide mass shifts are smaller. If the substrate is a small molecule (e.g. glucose), but contains multiple atoms per molecule of the element to be labeled, ideally only one of the atoms is labeled (or a small portion of atoms if it is a very large molecule) to avoid isotope “clumping”, as this can lead to a reduction in sensitivity (Fig. 3). Calis-p can, however, handle “clumped” data if needed. Similarly, if a complex substrate is used (e.g. complete plant leaves) ideally the complex substrate should only be partially labeled (e.g. by growing plants in an atmosphere with 10% of the CO_2_ being labeled) rather than using fully labeled substrate.

**Figure 8:**
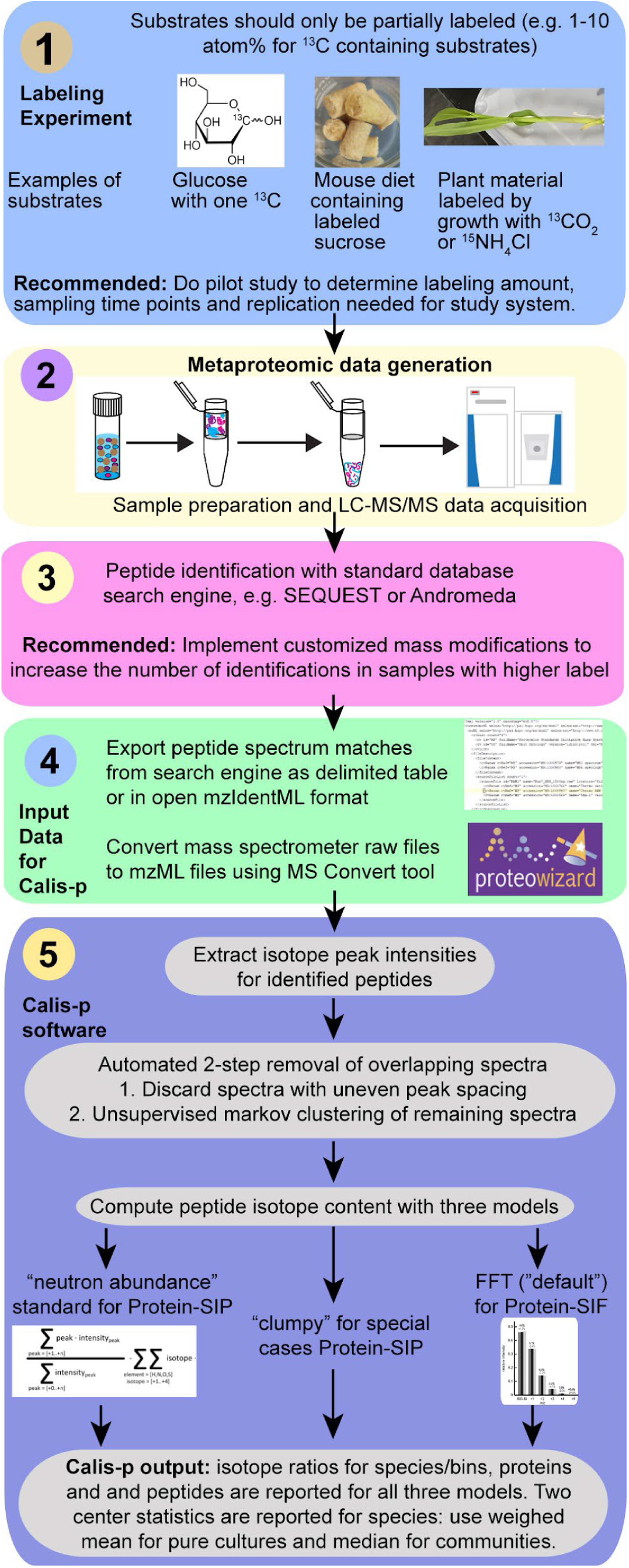
Protein-SIP and direct Protein-SIF workflow using Calis-p 2.1. The data filtering and computations illustrated in step (5) all happen in Calis-p in a fully automated fashion. The user has the ability to set specific parameters when starting the program. Full details on how to operate Calis-p are provided in the Wiki at https://sourceforge.net/projects/calis-p/. Not shown in the figure is that for Protein-SIF calibration of values with a reference material is needed, for details on this see the supplementary text and the original Protein-SIF publication^[14]^.

Other considerations for the labeling experiments include the number of replicates that are required, which depends on the biological question of the experiment, if a time course or a single time point will be sampled, and if a control with unlabeled substrate will be carried out, which is not needed for Calis-p, but can be helpful in data interpretation. Generally, we recommend to carry out a feasibility study, if at all possible, to determine the correct amount of label that works for the study system and time points that need to be sampled. Measurement of bulk label incorporation using an isotope ratio mass spectrometer can be useful in determining if an experiment worked prior to starting sample preparation for Protein-SIP.

The produced samples should be processed with a standard metaproteomic sample preparation method tuned to the particular sample type. In contrast to the protein-SIF method ^[14]^, which requires calibration for a small isotope offset caused by the instrument, no calibration reference material needs to be prepared for Protein-SIP. The produced peptide mixtures need to be analyzed by 1D or 2D liquid chromatography (LC) and tandem mass spectrometry (MS/MS) using a high-resolution Orbitrap mass spectrometer with standard metaproteomic LC-MS/MS approaches (see Methods and e.g. ^[41]^). One important consideration for the data acquisition in the mass spectrometer is the choice of resolution particularly for experiments involving ^15^N labeling (see Suppl. Results and Discussion).

The steps for data preparation for Calis-p and the computational steps implemented in Calis-p are described in detail in the Methods and on the Calis-p software repository website (https://sourceforge.net/projects/calis-p/).

## Discussion

The developed Protein-SIP approach provides a means to detect and quantify the incorporation of stable isotopes from labeled substrates into many individual species in microbial communities in one LC-MS/MS measurement and with minimal computational cost. Our approach has many advantages over other SIP approaches and previously developed Protein-SIP approaches. First, the approach allows for high throughput, as compared to most other stable isotope probing methods, such as DNA/RNA-SIP and nanoSIMS because as little as 2 hours of LC-MS/MS time will allow to quantify label incorporation for a good number of the more abundant species in a sample. For example, in bioreactors with 63 species we were consistently able to obtain sufficient measurement depth to quantify heavy water incorporation in >=20 species (Fig. 6). In contrast, nanoSIMSonly allows for measurement of isotope incorporation into a limited number of individual cells of very few species (2-3) in this time frame as species assignment of cells depends on species specific probes, and DNA-SIP is limited by the number of samples that fit into the ultracentrifuge rotor (usually six) needed for fine scale separation of heavy and light DNA and the cost associated with sequencing of a great number of individual density gradient fractions per sample. Second, our approach is a departure from previously developed Protein-SIP approaches in that it is highly sensitive and affords a large dynamic range of three orders of magnitude detecting label incorporation in the range of 0.01-10% of added label, while previous Protein-SIP detection principles require much higher label amounts to enable detection and usually offer only a dynamic range of one order of magnitude^[21]^. Similar is true for DNA/RNA-SIP based approaches, which require at least 20% label for detection^[8]^. The high sensitivity, accuracy and large dynamic range of our approach brings numerous advantages, including significant cost reduction due to lower use of often very expensive isotopically labeled substrates, the ability to work with much shorter labeling times, and simultaneous detection of label incorporation in slow and fast growing microorganisms. A massive cost reduction compared to other SIP approaches is possible, because most existing (Protein-)SIP approaches use upward of 20% (most often 100%) labeled substrate, while when using the Calis-p approach experiments with 1-10% label can be done cutting the experimental isotope use by 50-99%.

Using shorter incubation times is possible because incorporation of labels into proteins does not require for replication to occur, which is the case for DNA-SIP. It is important to note here that the ^13^C-label content of the substrate needs to be kept at 10% or below for our approach to work (higher percentages can be used for other elements see Box). A short labeling pulse with a substrate with higher label percentage would generate a heavy peptide population that would be completely mass shifted away from the unlabeled peptide population and thus become undetectable by Calis-p. Such strongly mass shifted peptide populations would be detectable with the MetaProSIP^[19]^ and SIPPER^[20]^ softwares. Third, we developed our approach to work with stable isotopes of all elements present in proteins, which allows tracking of assimilation of a large diversity of simple and complex substrates, as well as general activity markers such as ^18^O water. Based on our ^2^H and ^18^O case study results, we would recommend to use ^18^O water as the activity marker if compatible with the experimental design, as the current Calis-p version showed much higher sensitivity with the 18O data. More testing and optimization of Calis-p will be needed in the future for deuterated water using data to be generated with lower 2H labeling amounts. Fourth, Protein-SIP does not require isotope based separations of biological material such as the density gradient centrifugation used for DNA/RNA-SIP. That approach requires large amounts of material and sequencing of multiple fractions per sample. For this reason, Protein-SIP can be done with very small amounts of sample with an ideal starting amount of 1 mg or more of wet weight cell mass^[14]^. However, we have achieved good isotope estimates with as little as 50 μg using Calis-p for stable isotope fingerprinting^[42]^.

Currently, Protein-SIP only allows for labeling with one isotope per sample as changes in peptide isotope patterns cannot be attributed to specific elements. However, in the future it might be possible to develop Protein-SIP approaches that allow for parallel measurement of ^15^N and ^13^C incorporation in a single sample, because added neutron masses for ^15^N and ^13^C are sufficiently different from each other - due to differences in nuclear binding energy- to allow for their separation in ultra-high resolution mass spectrometers (Suppl. Results and Discussion). The current limitation for generating ultra-high resolution data suitable for separating peptide carbon and nitrogen isotopes is that higher resolution comes at slower mass spectrometric acquisition time. Thus, there is a tradeoff between ultra-high resolution data acquisition and obtaining a large number of MS^2^ spectra for peptide identification. Instruments with faster acquisition times and potentially alternative data acquisition modes such as data-independent acquisition (DIA) metaproteomics could make dual-label Protein-SIP feasible in the next few years.

## Methods

### Generation of labeled pure culture samples

The following steps were followed for single-carbon labeled and six-carbon labeled ^13^C glucose experiments with both *Escherichia coli* K12 (Obtained from Salmonella Genetic Stock Centre at the University of Calgary, Catalogue # SGSC 268) and *Bacillus subtilis* strain ATCC 6051. M9 and *Bacillus* minimal media were prepared without glucose. For M9 minimal medium we dissolved Na_2_HPO_4_ (12.8 g), KH_2_PO_4_ (3.0 g), NaCl (0.5 g), NH_4_Cl (1.0 g) in DI Water (978 ml) and autoclaved. Once the solution had cooled, we added the following filter-sterilized solutions: 1 M MgSO_4_ (2 ml), 1 M CaCl_2_ (0.1 ml), and 0.5% w/v thiamine (0.1 ml). Bacillus minimal medium (0.062 M K_2_HPO_4_, 0.044 M KH_2_PO_4,_ 0.015 M (NH_4_)_2_SO_4,_ 0.000 8 M MgSO_4_ x 7 H_2_O) was prepared, the pH adjusted to 7 and autoclaved. 20% stock solutions of both unlabeled and ^13^C-labeled glucose were combined to make a total of eight glucose mixes with final ^13^C-labeling percentages (% w/w) as follows: 0, 0.01, 0.025, 0.1, 0.25, 1, 5, and 10. Please note that the unlabeled glucose contained natural abundances of ^13^C of around 1.1% and that the percentage of ^13^C from labeled glucose has to be added to this. For unlabeled glucose we used D-(+)-glucose (> 99.5%) from Sigma Life Science, cat no. G7021 and for labeled glucose we used either D-glucose-U-^13^C (99%, Cambridge Isotope Laboratories, cat no CLM-1396-10) or D-glucose-2-^13^C glucose (99%, Aldrich, cat no. 310794). **Cell growth**: Frozen stock cultures were streaked on LB agar plates and incubated overnight at 37°C. A single colony was picked from the plate and grown overnight at 37°C in liquid media. Nine milliliters of overnight culture were spun down at 18,000 g for five minutes, the supernatant was discarded and pellets were washed twice with PBS to remove unlabeled glucose. Pellets were resuspended in 1 ml PBS. **Labeling**: Ten milliliters of liquid media without glucose were aliquoted into a total of 24 serum bottles per strain (triplicate bottles for each of the eight ^12^C/^13^C glucose mixes). 200 μl of the ^12^C/^13^C mixes and 10 μl of overnight culture were added into the serum bottles.

The bottles were then crimped, the headspace was flushed three times with CO_2_-free air and cultures were incubated overnight at 37°C while shaking at 100 RPM. **Sample processing**: Serum bottles were depressurized by inserting a sterilized needle into the septum to release air. Ten milliliters of culture from each bottle were spun down at 18,000 g for five minutes. The supernatant was discarded and the pellet resuspended in 2 ml of PBS to make two 1 ml aliquots. 50 μl of 1%, 5% and 10%-labeled glucose grown cells were used for cell counts using a Neubauer counting chamber. Cells were pelleted at 10,000 g for five minutes, the supernatant was discarded and pellets were flash-frozen in liquid nitrogen before being transferred to -80°C.

### Mock community spike-in experiments

The generation of the mock community (UNEVEN type) is described in Kleiner et al. (2017)^[22]^. We mixed *E. coli* cells grown in 1, 5 and 10% ^13^C_6_-labeled glucose containing media into three replicate samples of this mock community. We mixed the labeled *E. coli* cells in a 1:1 ratio to unlabeled *E. coli* cells already present in the mock community based on cell counts.

### Heavy water incubations of a microbial community derived from the human intestinal tract

The growth conditions, sample preparation and LC-MS/MS methods for the human intestinal microbiota grown in bioreactors has been described in Starke et al.^[11]^. Briefly, two bioreactors were inoculated with 63 bacterial strains (six phyla) isolated from a healthy human fecal sample. Bioreactors were fed with two custom media formulations representing different diets - high fiber and a high protein (see table S2 in ^[11]^). 2 ml batch cultures were set up using material from the bioreactors and 1 ml of pre-reduced, double strength medium (high fiber or protein), as well as 1 ml of unlabeled, ^18^O or ^2^H water was added. After a 12 h incubation at 37 °C in an anaerobic chamber samples were collected by centrifugation. The protein sequence database for identification of peptides from these samples was generated from the Uniprot reference proteomes for the species most closely related to the 63 isolates based on the 16S rRNA information published in Starke et al.^[11]^. When computing ^18^O and ^2^H abundances with Calis-p for these samples we corrected for offset, which can be caused by the natural deviation of the ^13^C abundance from the standard value as described in the supplementary methods.

### Sample preparation and One-Dimensional (1D) LC-MS/MS

Peptide samples for proteomics were prepared as described by Kleiner et al. (2017)^[22]^ following the filter-aided sample preparation protocol described by Wisniewski et al. (2009)^[43]^. Peptide concentrations were quantified using a Qubit® Protein Assay Kit (Thermo Fisher Scientific).

#### 1D-LC-MS/MS

Samples were analyzed by 1D-LC-MS/MS as described in Hinzke et al. (2019)^[41]^. Replicate samples (e.g. replicate 1 at 1%, 5% and 10%) were run consecutively followed by two wash runs and a blank run to reduce carryover. For 1D-LC-MS/MS, 0.4 μg (pure culture samples) or 2 μg of peptide (mock community-spike in samples) were loaded onto a 5 mm, 300 μm i.d. C18 Acclaim PepMap 100 precolumn (Thermo Fisher Scientific) using an UltiMate 3000 RSLCnano Liquid Chromatograph (Thermo Fisher Scientific). After loading, the precolumn was switched in line with either a 50 cm × 75 μm (pure culture samples) or a 75 cm × 75 μm (mock community – spike in samples) analytical EASY-Spray column packed with PepMap RSLC C_18_, 2 μm material. The analytical column was connected via an Easy-Spray source to a Q Exactive Plus hybrid quadrupole-Orbitrap mass spectrometer (Thermo Fisher Scientific). Peptides were separated on the analytical column using 140 (pure culture samples) or 260 (mock community - spike in) min gradients and mass spectra were acquired in the Orbitrap as described by Petersen et al. (2016). The resolution used on the Q Exactive Plus for MS^1^ scans, which provide the isotope pattern information used by Calis-p, was 70,000.

### Peptide identification and data preparation for Calis-p

Briefly, the LC-MS/MS data were used as the input for peptide identification using the database search engine SEQUEST HT implemented in Proteome Discoverer 2.2 (Thermo Scientific). Note, other standard search engines such as Andromeda implemented in MaxQuant^[44]^ can be used as well. We used experiments specific protein sequence databases for the searches and these databases have been submitted along with the LC-MS/MS data sets (see Data Availability). Taxonomic information available for protein sequences in the search database, for example from metagenomic binning and classification, was indicated as a prefix in the accession number (e.g. >TAX_00000) to enable Calis-p to report isotope values for each taxonomic group. The searches were modified to increase peptide identification rates for higher label amounts using customized modifications (see Suppl. Results and Discussion). The peptide spectrum matches (PSMs) produced by the search engine were exported from the search engine either in tabular format or in the open format mzIdentML and provided to Calis-p together with the mass spectrometry raw data in the open mzML format. The mzML files were generated from the raw data using MSConvertGUI via ProteoWizard^[45]^ with the following options set: Output format: mzML, Binary encoding precision: 64-bit, Write index: checked, TPP compatibility: checked, Filter: Peak Picking, Algorithm: Vendor, MS Levels: 1 (The MS/MS scans are not needed for isotope pattern extraction).

Once input files and optional parameters are provided Calis-p extracts isotope patterns for all identified peptides using a procedure optimized for Protein-SIP. The isotope patterns are extensively filtered for quality and high quality patterns are used for calculation of peptide isotope content using three different models. The “default” model developed for Protein-SIF, the “neutron abundance” model, which usually works best for Protein-SIP, and the “clumpy” model (see Methods). Calis-p automatically provides output files for all three models for taxa, proteins and peptides in a tabulated format that can subsequently be used in statistical and other data analysis softwares such as R.

### SIP computation algorithms and computational improvements to increase speed and accuracy of isotopic pattern extraction

As a starting point for estimation of stable isotope composition of isotopically labeled samples, we augmented the Calis-p software previously developed for estimation of ^13^C at natural abundance^[14]^. For estimating natural ^13^C abundance the software uses a model that assumes random distribution of ^13^C atoms in peptides, leading to peptide spectra with predictable isotope patterns. These isotope patterns are modelled in Calis-p with Fast Fourier Transformations (FFT). With labeled samples, the shape of spectra cannot be predicted using FFT, because these spectra become mixtures of spectra associated with labeled and unlabeled peptides. Both the proportion of heavy isotopes in the labeled peptides and the extent of labeling - the relative abundances of labeled versus unlabeled populations of peptides - are unknown in advance.. Therefore, we used the following more general equation to infer the number of neutrons from peptide isotope patterns to implement a “neutron abundance” model:

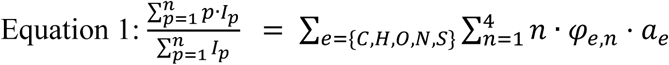

With, on the left, considering an isotope pattern of size *n* peaks, *p* is the peak number, and *I* is the intensity of peak *p*. On the right, for each isotope, *e* is its element [C,H,O,N,S], *n* the number of additional neutrons, *φ* its abundance (fraction), and *a* the number of atoms of the element in the peptide associated with the spectrum. Table 1 shows the estimates for natural abundances of the isotopes used in calculations.

**Table 1.** Estimates for natural abundances of the isotopes used in calculations based on the IUPAC Technical Report on the Atomic Weights of the Elements ^[46]^.

|  | Isotope |  |  |  |  |
| --- | --- | --- | --- | --- | --- |
| Element | +0 | +1 | +2 | +3 | +4 |
| C | 0.9889434148335 | 0.011056585 | 0 | 0 | 0 |
| N | 0.996323567 | 0.003676433 | 0 | 0 | 0 |
| O | 0.997574195 | 0.00038 | 0.002045805 | 0 | 0 |
| H | 0.99988 | 0.00012 | 0 | 0 | 0 |
| S | 0.9493 | 0.0076 | 0.0429 | 0 | 0.0002 |

We can rearrange this equation to, for example, calculate the fraction of ^13^C, assuming all other isotopes are at natural abundance, as follows:

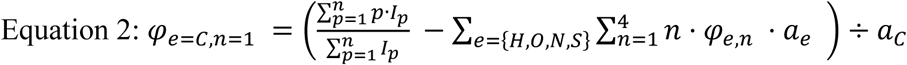

The second SIP computation algorithm was implemented in Calis-p as the “clumpy label” model. When labeling with substrates that contain multiple isotopically labeled atoms, for example fully labeled ^13^C_1-6_ glucose, this can lead to assimilation of clumps of labeled atoms into a single amino acid. For example, fully labeled glucose will be converted to fully labeled pyruvate, which, in turn, will be converted to fully labeled alanine, which will be incorporated into protein. This leads to peptide spectra that display higher-than-expected intensity at a higher isotopic peak numbers. To estimate the “clumpiness” of heavy isotopes in peptides, we developed the following procedure (detailed explanation in Figure S6): First, only the monoisotopic peak (A = +0) and A+1 peaks of the spectrum are used to estimate the fraction assimilated in clumps of one heavy atom (e.g. ^13^C). Next, the experimental intensity of the A+2 peak is compared to its expected intensity assuming all label was assimilated in clumps of one heavy atom. Any additional intensity of the A+2 peak is assigned to assimilation of clumps of two heavy atoms. This way, all peaks up to A+6 are inspected. The algorithm assumes the peptides are completely labeled, i.e. labeled to saturation. Usually, stable isotope probing experiments do not proceed that long, but doing so would enable determination of the number of labeled atoms in the substrate assimilated by each species via this procedure.

In typical proteomics data, tens to hundreds of MS1 spectra are collected for each detected peptide, at different elution times and mass over charge ratios. MS1 spectra can be crowded, especially for samples from more complex microbial communities. Unfortunately, overlap between isotopic patterns associated with different peptides can lead to overestimation of labeling. We have added new filtering routines, which remove such compromised isotopic patterns in two steps. First, any isotopic patterns with uneven spacing between peaks (which could indicate overlap with another spectrum) are discarded. On average, the peaks that form an isotopic pattern associated with a given peptide are separated by 1.002 Da, divided by the charge z of the peptide. If a pattern’s median peak spacing was <1.000/z or >1.004/z, or if the average sum of squares of the difference between the actual spacings and the median spacing was > 1×10-5, the entire isotopic pattern was discarded.

Next, remaining spectra are filtered out by unsupervised Markov clustering of all remaining spectra associated with a peptide^[47]^. The premise of this filtering approach is that clean spectra will be similar to each other, while spectra affected by noise are likely to be more different from each other. After filtering, all remaining spectra are truncated to the most common number of peaks, and spectra with fewer peaks are discarded. Spectra are then normalized to a total intensity of *1*, and an average (weighed by total spectral intensity) normalized spectrum was calculated for each peptide. The averages are weighed by intensity because high intensity spectra are more accurate and less noisy.

The normalized spectrum of each peptide is used to estimate the peptide’s isotopic composition using the original “Fast Fourier Transformations” based model (also called “default”), as well as the new “neutron abundance” (Equation 1) and “clumpy label” models.. For each species and protein in the sample, two center statistics are calculated based on all peptides associated with a species or protein: the median and the intensity-weighted average. The supplementary methods provide a detailed discussion of which center statistic to use when.

### Generating an additional label incorporation measure and Increasing peptide identification by using mass shift modifications in peptide identification searches

In addition to estimates based on MS1 spectra, we also estimated the degree of labeling based on the output of the search engine used for peptide identification. For this, we defined six custom post-translational modifications in the search engine that enable the dynamic addition of 1-6 neutrons to a peptide during the search. We tested multiple implementations of these dynamic modifications (Figs. 2 & S2, Suppl. Results & Discussion). Details on the implementation of the modifications in a search engine can be found in the Calis-p software documentation (https://sourceforge.net/p/calis-p/wiki/PSM%20files/).

### Other improvements of the Calis-p software

In addition to expanded functionality with regard to filtering of peptides and labeling, the software was also improved in many other ways: It now computes isotopic content of peptides with post-translational modifications and peptides containing sulfur peptides. It finds many more MS1 spectra for each peptide by searching for spectra at additional mass to charge ratios. Next to tab-delimited text PSM files exported from Proteome Discoverer, it now also parses open source mzidentml XML files (http://www.psidev.info/mzidentml). Finally, code efficiency improvements and implementation of multi-threading led to much faster computation, requiring less than one minute to process all spectra recorded during a 2 h run on a QExactive Plus Orbitrap mass spectrometer, using 10 threads. Source code and more details about algorithms and procedures can be found at http://sourceforge.net/projects/calis-p/.

### Comparison with existing Protein-SIP tools

To benchmark and compare Calis-p against existing Protein-SIP tools under optimal operation conditions, we invited developers/operators of Sipros ^[17]^, MetaProSIP ^[19]^ and SIPPER ^[20]^ to participate in a tool comparison. Drs. Sachsenberg (MetaProSIP) and Tolić (SIPPER) joined our effort. We were unfortunately not able to find an operator for Sipros and were also unable to get the tool to work on our own. For the comparison we used four raw files from the mock community spike-in experiments including unlabeled, 1% label, 5%, label, and 10% label in the spiked in *E. coli* and the corresponding protein sequence database. For processing with Calis-p we used the optimized settings from this study.

The SIPPER tool was developed at the Pacific Northwest National Laboratory and is available as an open source software written in C#. SIPPER requires an unlabeled incubation MS dataset as a reference for extracting target peptide IDs to which all stable isotope incubated datasets are compared. To generate peptide-spectrum matches to be used for isotope estimates the MSGF+ search algorithm ^[48]^ was used to search the unlabeled sample against the protein sequence database. The precursor mass tolerance was set to 20 ppm and oxidation of M and N-terminal acetylation were included as dynamic modifications and carbamidomethylation of C as static modification –, The search provided 112,580 identified target sequences (including contaminant IDs). The parameters for the SIPPER isotope calculation run included summing 7 precursor spectra around each target scan number, a 10 ppm tolerance for mass accuracy, 10% tolerance for normalized elution time, and filter confidence ID criteria outlined in the manuscript.

The MetaProSIP tool is integrated into the OpenMS open source software GUI^[49]^. An OpenMS workflow including a database search with Comet followed by the MetaProSIP tool was built according to the recommended parameters from the original publication with the minor modification that we activated the MetaProSIP setting to subtract the mono isotopic peak value. A reference (unlabeled) incubation was not needed to perform isotope calculations, because we expected part of the *E. coli* peptide population in the sample to be unlabeled due to the presence of unlabeled E. coli in the mock community thus all four data files were run through the workflow. The database search parameters were 10 ppm mass tolerance for precursor ions, up to two missed cleavages, carbamidomethylation of C as static modification, and oxidation of M as dynamic modification. Results were reported as relative isotope abundances (RIA) per ID. The MetaProSIP algorithm can split the RIA distributions of peptides with higher abundances into multiple isotope abundance clusters, leading to multiple reported RIAs per peptide. This happened on average for ∼9% of peptides in our datasets (0.3% in unlabeled, 6% in 1% labeled, 14% in 5% labeled, and 13% in 10% labeled). If the algorithm provided multiple RIAs for a peptide, we only used the highest RIA value in all downstream calculations.

To make the output data from both SIPPER and MetaProSIP comparable to the Calis-p output we used the tables from both approaches that report 13C content per PSM or peptide. SIPPER 13C content was reported in terms of percent carbon and percent peptides labeled, MetaProSIP 13C content was reported as relative isotope abundance, and Calis-p ^13^C/^12^C ratios were converted to 13C atom percent. We filtered the data to only include distinct protein unique peptides then calculated the 13C content for each species by taking the median. For visual representations (Fig. S4), we additionally filtered the data for which 13C values were provided for at least 9 peptides per species (equivalent to filter used for Calis-p).

### Data and software availability

The Calis-p (version 2.1) was implemented in Java and is freely available for download, use and modification at http://sourceforge.net/projects/calis-p/ and with a stable identifier at Zenodo https://doi.org/10.5281/zenodo.5619585 (Reviewer access can be obtained via this link https://datadryad.org/stash/share/HYoUsJAbcrkBaoPumrts4rcxSTMHO1iifjL-5YE3SU0).

The MS proteomics data and the protein sequence databases have been deposited to the ProteomeXchange Consortium ^[50]^ via the PRIDE partner repository with the following dataset identifiers: *B. subtilis* and *E. coli* grown in minimal medium with fully labeled ^13^C_1-6_-glucose at different concentrations PXD023693 [Reviewer Access at: https://www.ebi.ac.uk/pride/login User: Password: 0AyI3dGY], *B. subtilis* and *E. coli* grown in minimal medium with singly labeled ^13^C_2_-glucose at different concentrations PXD024285 [Reviewer Access at: https://www.ebi.ac.uk/pride/login User: Password: 0oSaTgAW], labeled E. coli spiked into the mock community PXD024174 [Reviewer Access at: https://www.ebi.ac.uk/pride/login User: Password: axKUZrEV], mixing of labeled and unlabeled *E. coli* PXD024287 [Reviewer Access at: https://www.ebi.ac.uk/pride/login User: Password: VmwP04XB], *E. coli* labeled to saturation with 2.5% ^15^N ammonium PXD024288 [Reviewer Access at: https://www.ebi.ac.uk/pride/login User: Password: oQcd2WjH], and the human microbiota derived microbial community from Starke et al. ^[11]^ grown with heavy water and different diets PXD024291 [Reviewer Access at: https://www.ebi.ac.uk/pride/login User: Password: GHbI87wI].

The mock community data without labeled *E. coli* spike-in was previously published ^[22]^ and we retrieved files Run4_U2_4600ng.msf and Run5_U2_4600ng.msf from PRIDE Project PXD006118.

## Author contributions

M.K. and M.S. designed research; M.K., A.Kouris., J.M., and M.S. performed research; M.K., A.Kouris., M.J., Y.L., A.Korenek, G.D., T.S., N.T., M.L., and M.S. analyzed data; and M.K. and M.S. wrote the paper with support from all co-authors.

## Acknowledgements

We thank Nico Jehmlich and Robert Starke for providing the raw data for the heavy water dataset, and J. Alfredo Blakely-Ruiz for comments on the manuscript.

This work was supported by USDA National Institute of Food and Agriculture Hatch project 1014212 (MK), by the National Institute Of General Medical Sciences of the National Institutes of Health under Award Number R35GM138362 (MK), the U.S. National Science Foundation grant OIA #1934844, the Novo Nordisk Foundation INTERACT project under Grant number NNF19SA0059360 (MK), the Foundation for Food and Agriculture Research Grant ID: 593607 (MK), the Canada Foundation for Innovation (#32181, MS), the Natural Sciences and Engineering Research Council (NSERC), the Canada First Research Excellence Fund (CFREF), the Government of Alberta, and the University of Calgary.

