## Supplemental Text and Figures for "Ultra-sensitive isotope probing to quantify activity and substrate assimilation in microbiomes"

Manuel Kleiner: manuel_kleiner (AT) ncsu.edu

Marc Strous: mstrous (AT) ucalgary.ca

### Supplementary Results and Discussion

#### Differences when using Calis-p for quantification of natural carbon isotope ratios (Protein-SIF) versus labeling with heavy isotopes (Protein-SIP)

The new Calis-p version did not only enable the estimation of label content in Protein-SIP experiments, but the improved algorithms did also improve the estimation of natural carbon isotope abundances further, as compared to the previous Calis-p version [1]. The improved filtering of spectra and spectral clustering (see methods) led to a significant speed up of computation of the Fast Fourier Transforms (FFT) as fewer spectra have to be considered and additionally led to a higher number of peptides that passed the post FFT goodness-of-fit filtering threshold (Tables S1 & S2).

To benchmark the Protein-SIF capabilities of the new Calis-p version we re-analyzed a large dataset of metaproteomic data generated from mock microbial communities (details in [1]) to estimate the natural carbon isotope content in individual species in the communities and compare them to their known isotope content determined with isotope ratio mass spectrometry (IRMS)(Fig. S1). The new Calis-p version led to a significant increase in accuracy, for example the average difference between the actual and estimated $\delta^{13}C$ values for the fifteen most abundant organisms of the mock community decreased from 3.6‰ in the old version to 2‰ in the new version (Table S2). One critical change that led to this improvement in addition to improved filtering is the introduction of the median as an alternative center statistic (see supplementary method section on center statistics). As mentioned the improved filtering also led to an increase in the amount of data available for SIF calculations. For example, across all datasets 4 species had >500 peptides available for SIF calculation in the previous Calis-p version, while it was 11 species for the new version (Fig. S1).

When analyzing Protein-SIF data versus Protein-SIP data there are some important differences to consider, including:

(1) Protein-SIF requires the measurement of a calibration standard (e.g. human hair) with a known carbon isotope ratio as described in [1]

(2) While Protein-SIP experiments can be done and analyzed with Calis-p for all elements contained in unmodified proteins (C, H, N, O, S), Protein-SIF only works for carbon isotope estimates and Calis-p should not be used to try estimating SIF values for other elements as the results will be nonsensical.

(3) While the new "neutron abundance" model works best for Protein-SIP and also provides reasonable estimates for Protein-SIF, the FFT ("default model") algorithm, continues to provide the highest accuracy for Protein-SIF and thus for Protein-SIF, this model should be used.

(4) Similar to Protein-SIP, we showed here that using the median instead of the mean is advantageous, as it provides a significant improvement in accuracy for microbial community data and thus the median should always be used with microbial communities.

(5) While for Protein-SIP as few as 9 peptides (post-filtering) are sufficient to detect the presence of label, at least around 30 peptides (or 100 PSMs) are needed to get an accurate estimate of SIF values (Fig. S1).
(6) While Protein-SIP allows for the detection of label incorporation in individual proteins, PSM numbers for individual proteins are usually insufficient to obtain SIF values for individual proteins and thus SIF estimates are generally restricted to species/strain level estimates.

| | Protein-SIF accuracy of the prevsious calis-p version (v0.0, Kleiner et al. 2018 PNAS) | Protein-SIF accuracy of the new calis-p version (v2.0, this study) |
|---|---|---|
| **Intensity weighed mean δ¹³C accuracy as function of peptide numbers post-filtering** | 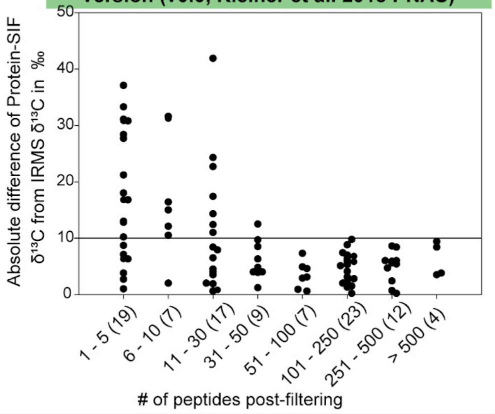 | 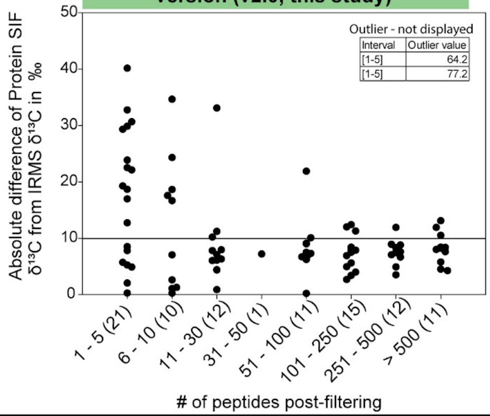 |
| **Median δ¹³C accuracy as function of peptide numbers post-filtering** | Not Available in v0.0 | 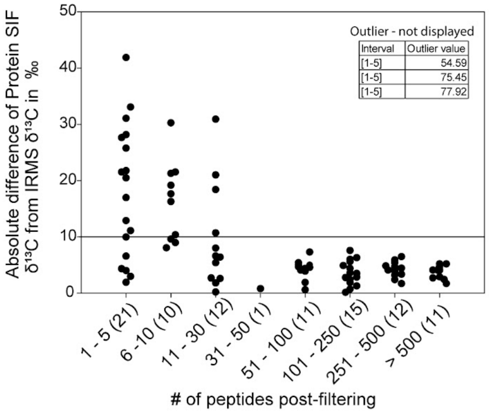 |
| **Intensity weighed mean δ¹³C accuracy as function of PSM numbers post-filtering** | 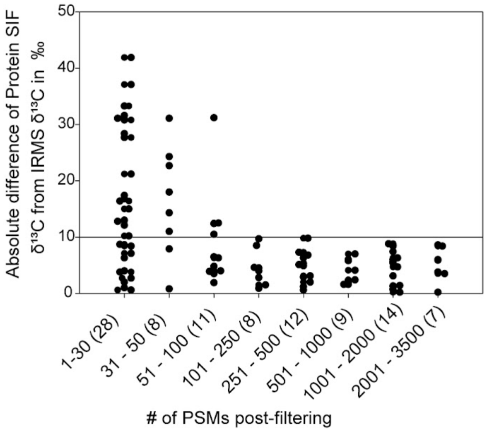 | 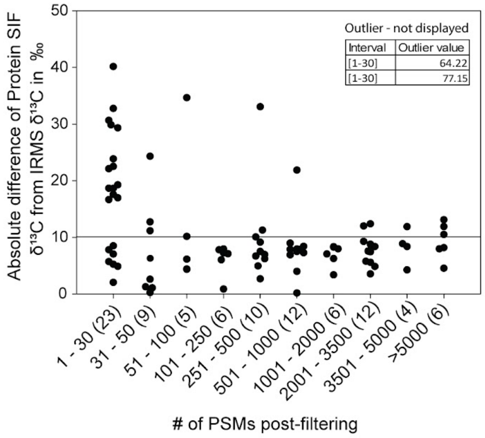 |
| **Median δ¹³C accuracy as function of PSM numbers post-filtering** | Not Available in v0.0 | 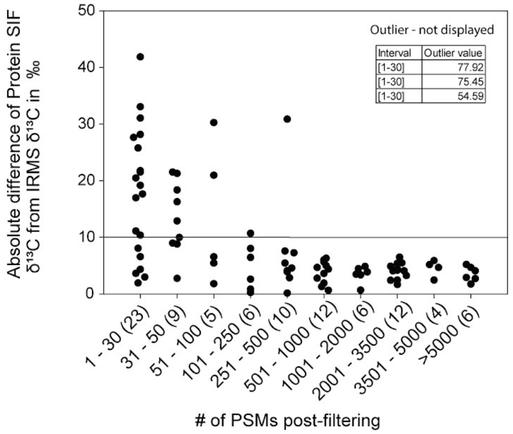 |

**Figure S1:** Comparison of the previously published version of Calis-p (v0.0)[1] with the new version (v2.1) in regards to their accuracy for quantifying natural carbon isotope abundances (Protein-SIF) of species in microbial community samples. The absolute difference between $\delta^{13}C$ values of individual species in mock communities determined with the Fast Fourier Transforms algorithm (i.e. default model) and isotope ratio mass spectrometry (IRMS) of the corresponding pure cultures is shown (method details in [1]). Five mock community datasets with a total of 32 species and strains were analyzed. For 20 species, the $\delta^{13}C$ values were known from IRMS performed on pure cultures. For these species, the $\delta^{13}C$ values were determined. Each dataset contained different amounts of data. The absolute difference between $\delta^{13}C$ values obtained via protein-SIF and IRMS was calculated and sorted according to how many peptides or PSMs were available for SIF calculation by Calis-p after filtering the peptides. The plots give the absolute differences for different ranges of peptide and PSM numbers used for SIF calculation. Additionally, for Calis-p v2.1 plots are also shown using the median as the center statistic for $\delta^{13}C$ value calculations.

#### A small modification of the peptide identification approach drastically increases the number of peptides with 1-10% label that can be identified

Before label content of peptides can be determined from their isotope patterns in the MS1 spectra, peptides need to be identified to obtain their sequence and sum formula. Peptide identification with standard identification algorithms such as SEQUEST[2], X! Tandem[3] or Andromeda[4] can be hampered at higher labeling percentages as they rely on the presence of the peptide's monoisotopic peak in the MS1 spectrum at natural isotope abundances, as well as peptide fragments at natural isotope abundance in the MS2 spectrum. With increasing label incorporation the intensity of the monoisotopic peak decreases and ultimately it disappears completely from the spectrum leading to loss of peptide identifications at higher label percentages (see Standard Search in Figs. 2 & S2).

To address this issue we tested two main strategies to allow for peptide identification with standard peptide identification tools even if isotope pattern shifts of variable size for individual peptides have occurred. The first strategy we used was an "Open Search"[5], in which the mass tolerance for the precursor (MS1) mass was set 20 Da instead of the usual values <0.1 Da. This strategy allows the peptide identification algorithm to consider mass shifts for the peptide mass of up to 20 Da, which increases the search space dramatically and thus computation time (e.g. 115x fold increase in search time as compared to our standard parameter search). For the second strategy we used four combinations of custom made dynamic modifications in the searches. These dynamic modifications, which are usually used to enable the identification algorithm to consider the presence of specific mass shifts due to post-translational modifications (PTMs) of peptides, increase the search space as well, but much less so as compared to the open search as only specific mass shifts are considered. The four combinations of dynamic modifications included: (1) "Dynamic Modifications" on all residues of a peptide with custom made 1 neutron and 2 neutron addition modifications allowing for up to three modifications of each per peptide. This results in up to 9 neutron mass additions in total per peptide. It is important to note that the exact mass shift to implement for a neutron depends on which element was labeled, because neutron mass in an element is influenced by the nuclear binding energy. For example, the exchange of one $^{14}N$ atom with one $^{15}N$ atom in a peptide changes its mass by 0.99703489 Da, while the exchange of one $^{12}C$ with one $^{13}C$ changes the mass by 1.0033548378 Da; (2-4)

Modifications on Termini: six custom modifications were set up that were restricted to either the C- or the N-terminus of the peptide and each of the modifications could only be present once. The modifications accounted for mass shifts of 1 to 6 neutrons and depending on the search strategy the low mass shifts (1, 2 and 3 neutron) were set up as modifications on the C- (N=high & C=low in Fig. 2) or the N-Terminus (C=high & N=low) or low and high mass shift modifications were distributed between both termini. Each modification could only be added to a terminus once. These strategies allowed for a total of 21 neutron additions to a peptide.

All strategies were tested on *Bacillus subtilis* ATCC 6051 and *Escherichia coli* K12 cultures that were labeled to completion using different percentages of $^{13}C_6$-glucose, as well as an unlabeled mock community consisting of 32 species [6] into which *E. coli* cells labeled at different percentages were spiked into (Fig. S2). As expected, peptide identifications decreased steeply at higher label percentages when using a standard identification strategy. For example, for the *B. subtilis* cultures average peptide-spectrum matches (PSMs) dropped from 12,888 at 0% added label to 1,483 at 10% $^{13}C$ label (Fig. 2a). The decrease was much less pronounced for *E. coli* PSMs in the spike-in samples (Fig. S2b), likely because the mock community contained also unlabeled *E. coli* cells in a 1:1 ratio to labeled cells allowing for peptide identification from unlabeled *E. coli* peptides. However, even in the spike-in samples a decrease of *E. coli* PSMs of more than 2-fold was observed between 1% and 10% added label. The Open Search strategy produced the lowest number of PSMs at all lower label percentages and performed slightly better than the Standard search at higher label percentages in some instances. However, for some of the Open searches on the mock community samples searches failed or we had to abort them after several weeks because they did not finish due to the magnitude of the search space. The fact that computation was already challenging for the relatively small sequence database used for the mock community disqualified this strategy as a viable approach. The Dynamic Modification strategy performed the best overall, particularly at higher label percentages, however, the searches for this strategy again failed in some instances or had to be aborted when searching the mock community data due to the high computation time needed. The three Modifications of Termini strategies performed equally well overall and consistently provided higher PSM numbers as compared to the Standard search, particularly at higher label percentages, outperforming the Standard search in some cases by several fold. The Modifications of Termini strategy with the low number of neutron additions on the N-terminus and the high number of neutron additions on the C-terminus performed slightly better than the other two strategies.

We chose to implement the Modifications of Termini (N=low, C=high) strategy in Calis-p as it provides a good balance between increase in PSMs obtained at higher labeling percentages and increase in computation time required. If the custom modifications are implemented in a search and correctly named (see https://sourceforge.net/p/calis-p/wiki/PSM%20files/) Calis-p will recognize them when reading the PSM table/mzIdentML file and count the number of neutron masses added by modifications to a peptide. The count is reported by Calis-p as the ratio of added neutrons divided by the total number of atoms in the peptide. While this count does not provide reliable quantitative information on label incorporation it does provide a line of evidence that the label was incorporated into a peptide/protein/taxon. In time series experiments, it can also show that the label amount is increasing or decreasing over time. In summary, using a search strategy with custom modifications has two advantages: (1) It increases the number of identified peptides (PSMs) when higher label amounts are used, which allows for more precise labeling estimates for proteins and taxa in Calis-p; and (2) the counting of neutrons added by the custom modifications can provide another line of evidence that label was incorporated, in addition to the MS1 spectrum based approaches implemented in Calis-p.

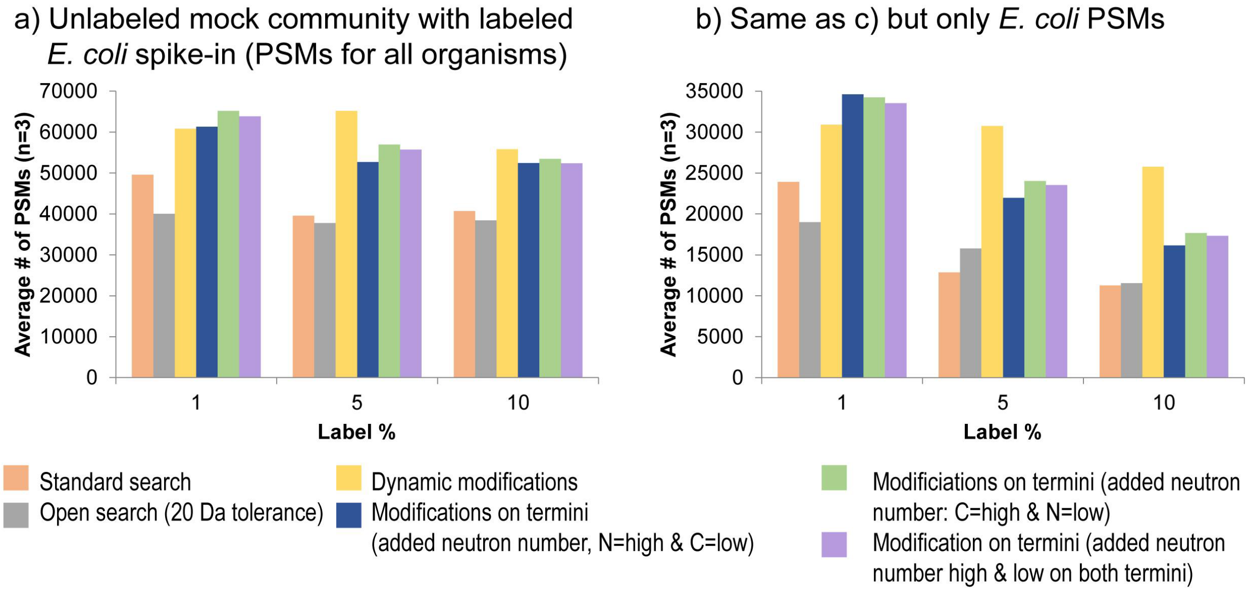


**Figure S2:** Number of peptide spectral matches (PSMs) identified at different $^{13}C$ label percentages using six different peptide identification strategies. For the spike-in samples *E. coli* cells labeled at different percentages with $^{13}C_6$ glucose were mixed into a mock community consisting of 32 species of bacteria, archaea, eukaryota and bacteriophages (UNEVEN community from Kleiner et al. (2017)[6]), which also contained unlabeled *E. coli* cells. Labeled and unlabeled *E. coli* cells in the spike-in sample were at a 1:1 ratio.Three biological replicates were analyzed for each label percentage. Peptides were identified using the SEQUEST HT Node in Proteome Discoverer (version 2.2.) with six different strategies to account for the mass shifts caused by addition of heavy atoms. Standard search: no dynamic modifications to account for addition of label; Open search: the precursor mass tolerance was set to 20 Da allowing for the potential addition of 20 neutrons (e.g. $^{13}C$ atoms) in a peptide; Dynamic modifications: allowing for up to three dynamic modifications each of two custom peptide modifications adding a 1 neutron mass shift and a 2 neutron mass shift (up to 9 neutrons in total per peptide); Modifications on termini: six dynamic modifications were set up that were restricted to either the C or the N-terminus of the peptide. The modifications account for mass shifts of 1 to 6 neutrons and depending on the search strategy the low mass shifts (1, 2 and 3 neutrons) were set up as modifications on the C or the N-Terminus or low and high mass shift modifications were distributed between both termini. Each modification can only be added to a terminus once. This strategy allows for a total of 21 neutron additions to a peptide.

#### Limit of detection in pure cultures of B. subtilis and E. coli

We used the data presented in Figure 3 and Suppl. Table S3 to estimate the limit of detection (LOD) of our method. For this we used a commonly accepted approach that gives the LOD as three standard deviations above the mean value of the baseline or control samples (see calculations below). The LODs were very low (13C/12C ratios of 0.01019 to 0.01026) indicating a labeling LOD of <0.01% with the exception of the B. subtilis samples labeled with 13C1-6 glucose for which the LOD 13C/12C ratio was 0.01055 indicating a labeling LOD of <0.1%.
To test if we could statistically differentiate labeled from unlabeled samples as close as possible to the LOD we used Student's t-test to compare the unlabeled with the labeled samples nearest the LOD. All resulting p-values were <0.01 indicating strong statistical support (see below).

Limit of detection (LOD) calculations

| | E. coli 13C2 | E. coli 13C1-6 | B. subtilis 13C2 | B. subtilis 13C1-6 |
|---|---|---|---|---|
| Mean of median 13C/12C ratios of 0% label samples | 0.01021 | 0.01017 | 0.01017 | 0.01025 |
| Standard Deviation | 0.000017 | 0.000005 | 0.000021 | 0.0001 |
| LOD | 0.01026 | 0.01019 | 0.01023 | 0.01055 |
| Label % at which all measured 13C/12C ratios are above LOD | 0.01 | 0.01 | 0.01 | 0.1 |
| T-test p-value of comparison of 13C/12C ratios for 0% labeled samples with samples at LOD | 0.0017 | 0.0018 | 0.005 | 0.00015 |

#### Mixing of labeled and unlabeled *E. coli* cells

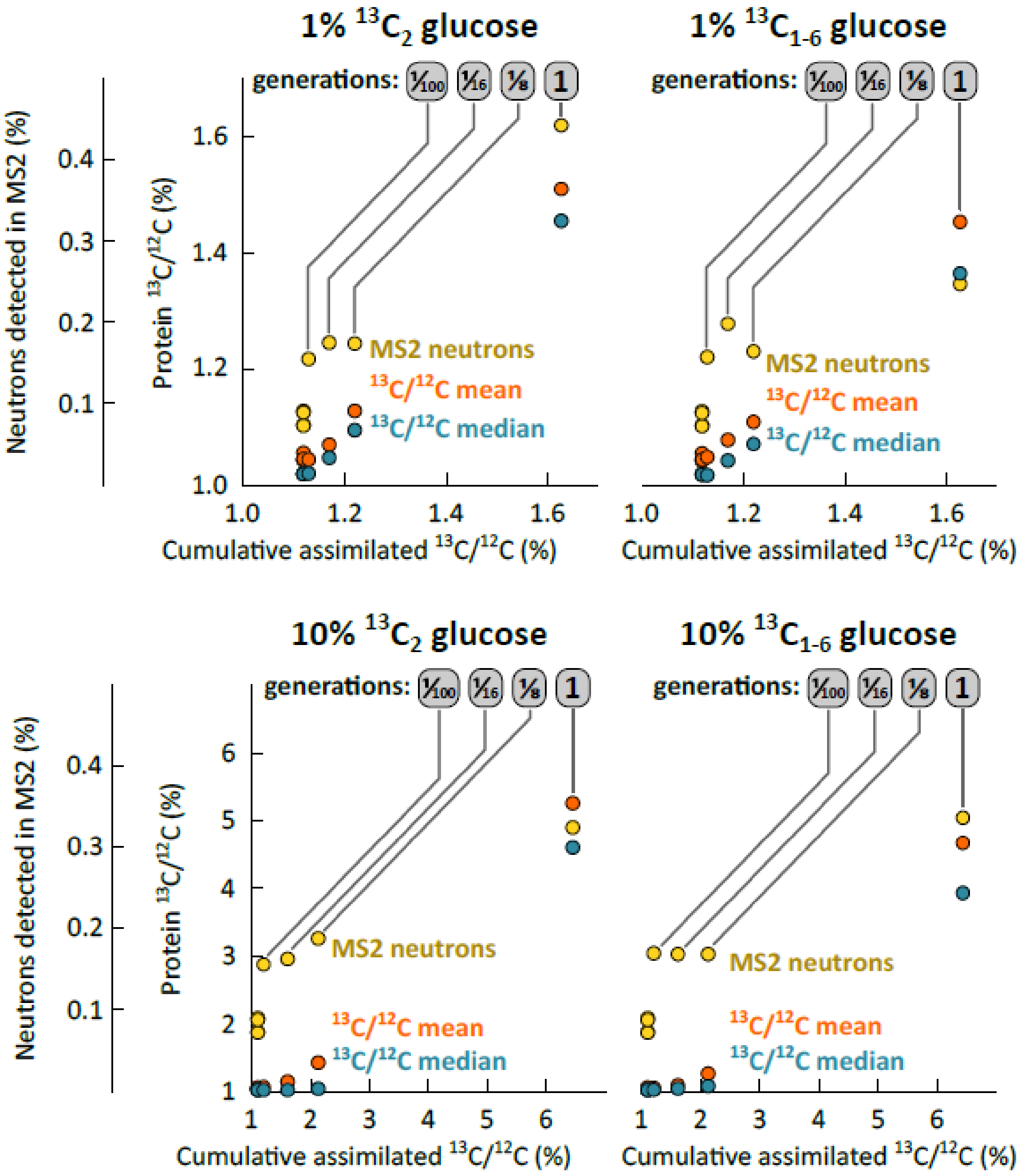


**Figure S3.** Assimilation of $^{13}C$ in a mock labeling experiment created by mixing labeled and unlabeled cells of *E. coli* in ratios corresponding to 1/100 to 1 generations of growth. Cells were labeled with 1% $^{13}C$ glucose (top row) and 10% $^{13}C$ glucose (bottom row). Cells were labeled with $C_2$-labeled glucose (left column) and fully ($C_{1\text{-}6}$) labeled glucose. Peptide identification based detection of excess neutron masses was most sensitive, able to reproducibly detect growth for 1/100 generation, but not quantitative. Use of 1% $^{13}C$ glucose with a single atom labeled resulted in the best quantitation ($R^2$ >0.996) and label recovery (92%). The detailed data for this figure can be found in Suppl. table S4.

#### Benchmarking with mock communities and spike-ins including detection limit and accuracy

To determine at what labeling level and at what organisms abundance within a microbial community we can still detect label incorporation into an organism and individual proteins we used a mock community. The mock community was previously established [6] and consisted of 32 bacteria, archaea, eukaryotes and viruses. Each of the species in the mock community was mixed in at a different abundance and the mock community consisted of abundant organisms (>5% abundance in terms of proteinaceous biomass) and low abundant organisms (<0.5%). In this mock community around half of the added *Escherichia coli* cells were labeled using $^{13}C_{1-6}$-Glucose, corresponding to one generation of labeling, with glucose containing 0, 1, 5, and 10% $^{13}C$. We generated triplicate samples for each labeling percentage.

For *E. coli* on average 574 peptides passed the quality filters in Calis-p and were used for calculation of the $^{13}C/^{12}C$ ratio. The measured ratios closely matched with the expected ratios and followed a linear trend with increasing label percentage (Fig. 4a and Table S5). Label incorporation was consistently detected down to 1% added label.

To test if label incorporation would also be detectable for low abundance organisms in the mock community, we downsampled the *E. coli* data to match the number of peptides passing the Calis-p quality filter for five lower abundant organisms (Fig. 4b to 4f). Additionally, *E. coli* peptides were chosen to match the MS1 intensity of peptides from the lower abundant organisms. We found that label incorporation could be detected with as little as one peptide, which corresponded to the number of peptides passing the Calis-p quality filters for *Nitrosomonas europaea,* which was the lowest abundance organism in the mock community making up 0.1% of the total proteinaceous biomass in the community (Fig. 4f). While detection of label incorporation was still feasible with very few peptides the accuracy of the measured ratios was low.

#### Choosing the right mass spectrometric resolution

It is important to note that in this study mass spectrometer resolution was set at 60,000 (120,000 for the heavy water data) and that it is critical to consider resolution depending on experiment type and labeled element used. At a resolution level of ~70,000 or lower isotopic peaks caused by neutron additions to different elements are not separated. Since neutron masses are different for isotopes of different elements due to differences in nuclear binding energy between elements (e.g. neutron mass is 1.0034 Da for C and 0.997 Da for N), at high enough resolution peptide isotope peaks caused by isotopes of different elements (e.g. $^{15}N$ and $^{13}C$) will be separated in the spectrum. For example, for a m/z 400 Da peptide a resolution of >60,000 will lead to separation of carbon and nitrogen isotope peaks, while hydrogen and oxygen isotope peaks will not be separated from carbon at this resolution. The carbon and nitrogen isotope peak separation impacts overall isotope peak intensities and therefore influences isotope abundance estimates by the three models implemented in Calis-p. Depending on which element is analyzed the impact of isotope separation in ultra-high resolution data (>100,000) will be different. For carbon we only expect this to impact natural abundance estimates with Protein-SIF and very low label percentage Protein-SIP data (<0.01% added $^{13}C$) due to the abundance of carbon atoms in peptides and the high natural abundance of $^{13}C$. For labeling experiments with deuterium and $^{18}O$ we also only expect impacts at very low labeling percentages because hydrogen, oxygen and carbon isotopes would not be separated at any reasonably achievable resolution and thus only the minor nitrogen isotope peak would be separated and impact peak intensities. However, for nitrogen labeled samples ultra-high resolution data will impact label calculations heavily as separation from the abundant carbon isotope peaks basically leads to a major subtraction of intensity. This scenario is currently not considered in the implemented models in Calis-p and therefore we recommend to only use resolutions of 70,000 or below for nitrogen labeling experiments. Additionally, for Protein-SIF and work with ultra-low $^{13}C$ label amounts we also recommend to remain below 70,000 resolution.

#### Comparison with existing Protein-SIP approaches

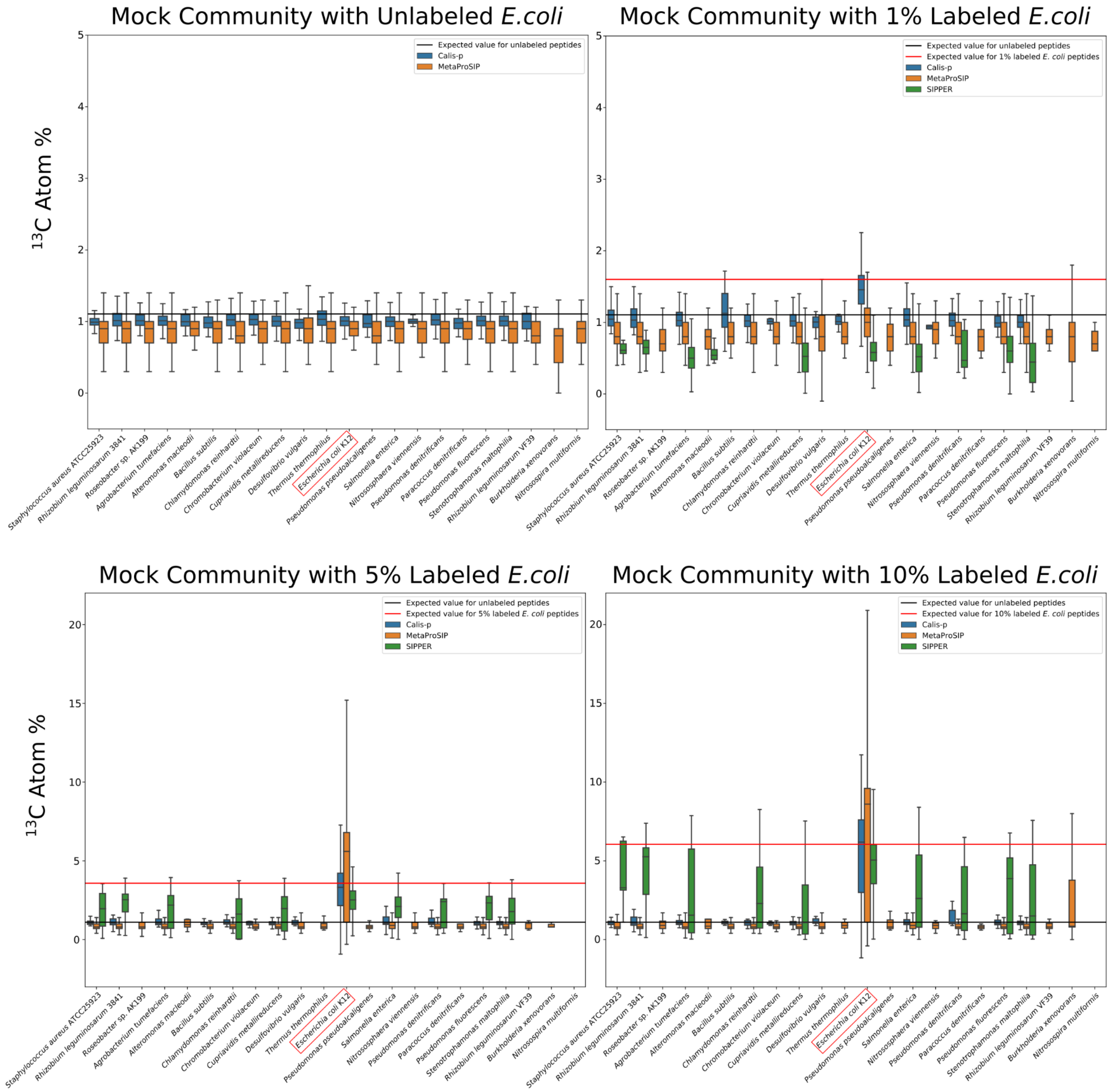


**Figure S4: Comparison of existing Protein-SIP approaches with Calis-p.** Three Protein-SIP approaches (SIPPER, MetaProSIP, and Calis-p) were compared using optimal parameters chosen by an expert operator for each tool. We used the datasets from the mock community with added 1%, 5%, or 10% 13C labeled E.coli and as a control the mock community with only unlabeled E. coli. The resulting $^{13}$C atom % output values from SIPPER, MetaProSIP, and Calis-p for each dataset were filtered such that organisms for which $^{13}$C content was quantified for 9 or more peptides were considered for plotting. The black line in each plot represents the expected $^{13}$C atom % value for all unlabeled peptides. The red line in each plot represents the expected $^{13}$C atom % value for the E. coli peptides from each respective dataset after accounting for the expected mixing of labeled and unlabeled E. coli peptides in the sample. The number of peptides used for each box and other details can be found in tables S7-S10.

#### Case Study: Differential heavy water incorporation in a bioreactor grown intestinal microbiota in response to high fiber or high protein

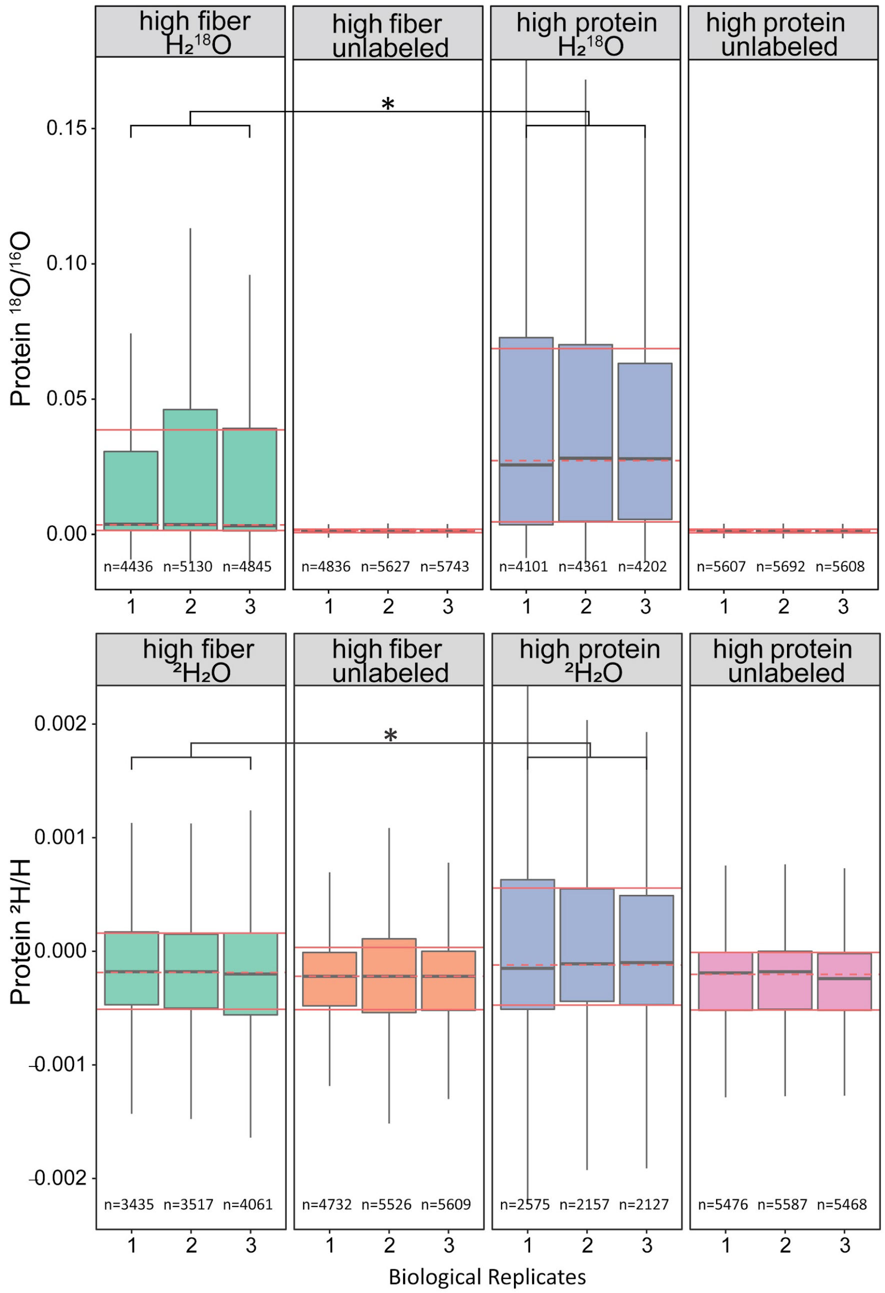


**Figure S5: Strong differences in heavy water incorporation in intestinal microbiota in response to different diet conditions.** 63 species isolated from human intestinal microbiota were grown together in triplicates in either a high fiber or high protein medium in the presence of unlabeled water or water with either 25% $^2H$ or $^{18}O$[7]. Calis-p based stable isotope ratios are shown for all species combined per replicate. Each box shows the data for all peptides for one replicate. The 'n' gives the number of peptides that passed the Calis-p quality filters. The solid red lines indicate the average quartiles for the three replicates, the dashed red line the average median for the three replicates. Statistically significant differences are indicated with '*' based on Student's t-test on the medians of replicates at $p<0.05$. The script (Suppl. File S1, also available here https://github.com/yihualiued/Data-analysis-of-Calis-p-protein-SIP-results/blob/main/Heavy%20Water_SIP_Calisp_final.R) and Calis-p output peptide datasets (Suppl. Datasets S1 and S2) used to generate this figure and Figure 6 have been provided as an example.

### Supplementary Methods

#### How Calis-p calculates center statistics for a species based on its peptides

Calis-p computes median and weighted mean estimators for $\delta^{13}C$ and $^{13}C/^{12}C$ ratios (or other heavy isotopes as instructed) for each species/genome/bin/MAG, each protein, as well as for each peptide spectrum match file and each spectrum file as a whole. These estimators are calculated separately for each of the three models. The means are weighed by intensity.
When using Calis-p with a pure culture, it makes sense that very intense peptides get more weight, because they are likely less affected by noise, so we recommend to use the weighted mean. However, for a mixed community, using the mean does not make sense because estimates of low-intensity peptides associated with a rare organism may be totally wrong when the spectrum has some overlap with a very intense spectrum associated with a more abundant organism. Therefore, for a mixed community it is better to use the median instead of the weighted mean.
Both center statistics are also calculated for each protein. Calis-p also performs a multiple regression to estimate weighted mean $\delta^{13}C$ (or other isotopes, as specified) values for each amino acid, again for all three models, organisms and files separately. In SIP experiments with labeled amino acids, this can show whether the labeled amino acid is taken up directly or indirectly, after the labeled amino acid was degraded by another species.
Instead of using center statistics produced by Calis-p, researchers can also find isotope estimates of all three models for individual peptides in the file peptides.csv. They could perform more sophisticated statistical analyses, tests and visualizations using data from that file, for example using R. Details on these files can be found on the Calis-p Wiki (https://sourceforge.net/p/calis-p/wiki/Center%20statistics/).

#### Correction of the used $^{13}C$ natural abundance value for calculation of other elements' isotope ratios

Since natural abundances of N, H, and O isotopes are very low, small natural differences in $^{13}C$ abundance can have a strong impact on calculated N, H, and O isotope ratios if an incorrect value for $^{13}C$ abundance is used. To allow us to correct for $^{13}C/^{12}C$ ratios that deviate from the expectation we implemented the natural isotope abundance table (Table 1) used in Calis-p for calculations with the "neutron abundance" model in a way that it can be updated with corrected values. For the default table we used published standard values from the IUPAC Technical Report on the Atomic Weights of the Elements [8]. This table can be exported (-

writeIsotopeMatrixFile ‘FILENAME’), corrected by changing natural abundance values, and then imported when executing Calis-p on a dataset (-readIsotopeMatrixFile ‘CORRECTEDFILENAME’).

We used this correction for the heavy water data from the intestinal microbiota bioreactors. For this, we first determined the $^{13}C$ abundance in unlabeled incubation samples computed by the Calis-p “default” model. We then used this $^{13}C$ abundance to correct the isotope abundance table for calculating the $^{2}H$ and $^{18}O$ abundances in the labeled samples.

#### Details for the procedure to estimate label "clumpiness"

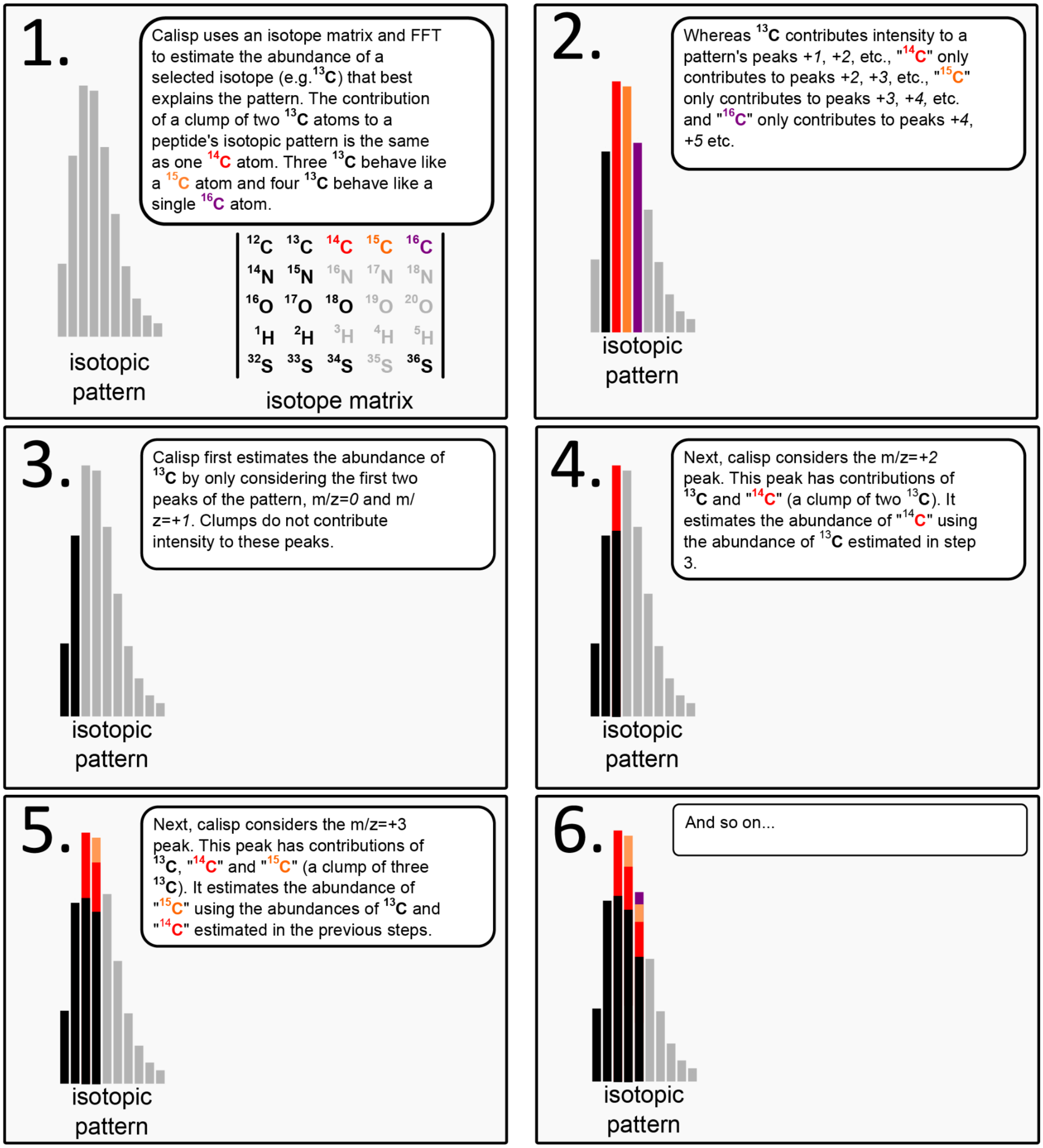


**Figure S6: Explanation of procedure how clumpiness of label is estimated.**
