## Supplementary material for "Ultra-sensitive isotope probing to quantify activity and substrate assimilation in microbiomes": File S1

# Yihua Liu, 2021-03-24
# Inputs files: 
#   - Calis-p peptides result: peptides.csv
#   - Taxonomy file: taxo.csv
#   - Parameters listed in the "Initialize WORKING DIRECTORY" section
# Targets: 
#   1. Data wrangling of "peptides.csv"
#   2. Visualize the isotope ratio distribution of peptides from different experiment groups
#   3. Statistical tests
#       
# ====== LOADING Libraries ======
library(tidyverse)
library(data.table)
library(ggstance)
library(Hmisc)

# ====== Initialize WORKING DIRECTORY ======

# Specify parameters (personalized aims)

# pepfile = "2H_SIF_Corrected_peptides.csv"             # specify the calis-p peptide file name/directory
# la = "H"                                              # specify the isotope label, "H" for D and "O" for 18O
pepfile = "18O_SIF_Corrected_peptides.csv"
la = "O"
isolim = "0.25"                                         # specify the isotope ratio cutoff for Calis-p, defined by the isotope ratio of labelling chemicals, as biomass cannot get higher ratio than label.
taxopath = "taxo.csv"                                   # specify the path of file containing Bin taxonomy info (relative to the path of this script)


# create a corresponding directory and move to the directory
# also assign texts to variables that specific to the label
if (la == "H" ){
  dir.create(str_c("2H", "Calis-p_Rslt_Prcs", Sys.Date(), sep = "_"))
  setwd(str_c("2H", "Calis-p_Rslt_Prcs", Sys.Date(), sep = "_"))
  lb <- "2H/H"
  lh <- "D2O"
}else if (la == "O"){
  dir.create(str_c("18O", "Calis-p_Rslt_Prcs", Sys.Date(), sep = "_"))
  setwd(str_c("18O", "Calis-p_Rslt_Prcs", Sys.Date(), sep = "_"))
  lb <- "18O/16O"
  lh <- "H218O"
}else {
  stop(cat("Error: Unspecified Isotope Label"))
}
   
       

# ====== Section 1. Data wrangling of "peptides.csv" ======
# Script to read in and cleanup peptide.csv
# Main process of data
#   1. clean up column names (merge the duo row column to one row)
#   2. remove cols from Default Model	and Clumpy Carbon Model (only use neutron abundance model)
#   3. remove peptides that matches more than one bins 
#   4. remove summed intensity = 0
#   5. replace peptide filename 


# Build a table to link sample numbers to corresponding experimental attributes 
samp_label <- tibble(
  sample_ID = 1:20,
  diets = c("fiber", "protein","fiber","fiber","fiber","protein","protein","protein","fiber","fiber","fiber","protein","protein","protein","fiber","fiber","fiber","protein","protein","protein"),
  label = c("Blank","Blank","unlabeled","unlabeled","unlabeled","unlabeled","unlabeled","unlabeled","D2O","D2O","D2O","D2O","D2O","D2O","H218O","H218O","H218O","H218O","H218O","H218O"),
  Rep = c(1,2,1,2,3,1,2,3,1,2,3,1,2,3,1,2,3,1,2,3)
) 
samp_label$sample_ID <- as.character(samp_label$sample_ID)


#read in peptides.csv data. Use designate col names to solve the incomplete first line.
#Warning message will appear as the rows are different in lengths (due to NAs at the end of rows)
pep <- read_tsv(str_c("../",pepfile), col_names = paste("X", 1:33, sep = "_")) %>%
  
  # Cut unnecessary columns (only use neutron abundance model)
  select(., -(X_15:X_24), -(X_28:X_33)) %>%
  
  # Assign actual col names from row1 and row2, 
  rename(.,
         Bin = X_1,
         Peptide_File = X_2,
         Spectrum_File = X_3,
         Protein = X_4,
         Peptide = X_5,
         Time_needed = X_6,
         Modifications = X_7,
         Flags = X_8,
         Elemental_composition = X_9,
         Mono_isotopic_mass = X_10,
         Neutrons_detected = X_11,
         Num_PSMs = X_12,
         Num_MS1_spectra = X_13,
         SI = X_14,
         Neutron_Abundance_Delta = X_25,
         Neutron_Abundance_Ratio = X_26,
         Neutron_Abundance_Spectrum = X_27) %>%
  
  # Cut first 2 rows (initial col names)
  # ---Till now, the two-line col names merged into one line
  filter(., Bin != "#" & Bin != "#Organism") %>%
  
  # Filter out rows with Flags (i.e. Too few spectra, No valid PSM, No majority vote while clustering, Too few spectra or Assigned to >1 orgamisms)
  # Note than this will not removing peptides matching one bin but multiple peptides
  filter(., is.na(Flags)) %>%
  select(., -(Flags)) %>%
  # There should be no NAs in Ratio col. But filter again to make sure
  filter(., Neutron_Abundance_Ratio == Neutron_Abundance_Ratio) %>%
  
  #Filter out crapped bins
  filter(., !str_starts(Protein,"CRAP")) %>%
  
  # Remove spectrum_File and extract sample_ID from peptides_File
  separate(Peptide_File, sep = "_", into = c("d1","d2","d3", "sample_ID", "d5","d6"))%>%
  select(-(d1:d3),-(d5:Spectrum_File), -(Time_needed:Modifications)) %>%
  
  # link sample ID with label and diet
  inner_join(.,samp_label, by = "sample_ID") %>%
  
  # build a col of group names
  mutate(expgp = str_c("high_", diets, "_", label))


# reading taxo info 
taxo <- read_csv(str_c("../",taxopath))


# Subsection: report the organisms with >9 peptides in each sample
# {
if (la == "H") {
  sampf <- samp_label %>%
    filter(label != "H218O")
  
} else {
  sampf <- samp_label %>%
    filter(label != "D2O")
  
}
organ_9pep <- tibble(x = c("number of organisms","sample ID"))
dir.create("organism count")
for (i in seq_along(sampf$sample_ID)) {
  organcount <- pep %>%
    filter(sample_ID == i) %>% 
    group_by(Bin)%>%
    summarise(PepNum = n())%>%
    filter(PepNum > 9) %>%
    # add taxo info
    inner_join(taxo, by = "Bin")
  write_csv(organcount, str_c("organism count/",la,"_organism_list_sample_",i,"_",Sys.Date(),".csv"))
  orgnum <- organcount %>%
    summarise(OrgNum = n())
  organ_9pep <- bind_cols(organ_9pep, tibble(x = c(orgnum$OrgNum,i)))
}
 write_csv(organ_9pep, str_c("organism count/",la,"_summary_organism_number_",Sys.Date(),".csv"))

 remove(organ_9pep,organcount, orgnum, i,sampf)
# }

# # filter out organisms that have less than 9 peptides in any sample
# # Creating a list of organisms that fit the criteria
 
OrganList <- pep %>%
  group_by(Bin, sample_ID) %>%
  summarise(PepNum = n()) %>%
  filter(PepNum >= 9) %>%
  # check the number of samples the organism presents
  ungroup() %>%
  group_by(Bin) %>%
  summarise(SampNum = n()) %>%
  filter(SampNum == 20) %>%
  select(Bin) %>%
  # add taxo info
  inner_join(taxo, by = "Bin")


# filter out entries with unrelevant labels
# also make a text label to print on plots and a text corresponds to "label" col
if (la == "H") {
  pep <- pep %>%
    filter(label != "H218O" & label != "Blank")

} else {
  pep <- pep %>%
    filter(label != "D2O" & label != "Blank")

}


# write file
write_csv(pep, str_c(la, "_peptides_Tidy_HeavyWater_" , Sys.Date(),".csv"))

# prepare a simplified data set for plotting
pepplot <- pep %>%
  mutate(Ratio = Neutron_Abundance_Ratio) %>%
  select(Bin:SI, Ratio, diets:expgp)%>%
  # coerce to data frame to avoid some problems
  as.data.frame() %>%
  # coerce SI and Ratio to numeric to avoid some problems
  mutate(SI = as.numeric(SI), Ratio  = as.numeric(Ratio)) %>%
  # remove points with isotope ratio goes over the limit, defined by label added.
  filter(., Ratio <= isolim)
# write file
write_csv(pepplot, str_c(la, "_peptides_Short_HeavyWater_" , Sys.Date(),".csv"))

remove(pep, taxo) 

# ====== Section 2. Visualize the isotope ratio distribution of peptides from different experiment groups =====
# That is, high protein unlabel vs high fiber unlabel vs high protein H218O vs high fiber H218O 
# Or, high protein unlabel vs high fiber unlabel vs high protein D2O vs high fiber D2O

# Box plots and statistics ####
# create a directory for box plots 
dir.create("Box plots")
# calculate data stats: the min, max and three quantiles of peptide isotopes

# for high fiber group and high protien group
stats_expgroups0 <- pepplot %>%
  group_by(expgp, Rep)%>%
  summarise(minRo = min(Ratio),
            y25 = wtd.quantile(Ratio, weights = SI, probs = 0.25),
            y50 = wtd.quantile(Ratio, weights = SI, probs = 0.5),
            y75 = wtd.quantile(Ratio, weights = SI, probs = 0.75),
            maxRo = max(Ratio),
            pepNum = n()) %>%
  # Note: in geom_boxplot, ymin is not the min number of y, but  
  #  "smallest observation greater than or equal to lower hinge (q1) - 1.5 * IQR"
  # where IQR ( inter-quartile range) = q3-q1
  # here instead, the following code use min of y or q1 - 1.5*IQR, whichever greater
  # similarly, ymax = max of y or q3 + 1.5*IQR, whichever smaller
  mutate(yIQR = y75 - y25) %>%
  mutate(y0 = ifelse(y25 - 1.5 * yIQR > minRo, y25 - 1.5 * yIQR, minRo),
         y100 = ifelse(y75 + 1.5 * yIQR < maxRo, y75 + 1.5 * yIQR, maxRo))
  
# calculate the quantiles' average for each exp group, as reference lines to show on the diagram
gpave <- stats_expgroups0 %>%
  group_by(expgp) %>%
  summarise(y25ave = mean(y25),
            y50ave = mean(y50),
            y75ave = mean(y75))

stats_expgroups <- inner_join(stats_expgroups0, gpave, by = "expgp") 

# Export boxplot data 
write.csv(stats_expgroups, str_c("./Box plots/",la, "_box_plot_exp_groups_",Sys.Date(),".csv"))


# Student's t-test for significance ----
 # select data to test: test the medium of 3 replicates from high protein against high fiber
 stat_gp <- stats_expgroups %>%
  filter(expgp %in% c(str_c("high_fiber_", lh),str_c("high_protein_", lh))) %>%
  separate(expgp,sep = "_", into = c("high","diets","label")) %>%
  select(Rep,diets,y50) %>%
  # spread the table
  spread(.,diets, y50) 
 # t-test
 t_gp <- capture.output(t.test(stat_gp$fiber, stat_gp$protein))
 # write t-test results
 write_lines(t_gp, str_c("./Box plots/",la,"_group_t-test_",Sys.Date(),".txt"))


# box plot for comparison across exp groups ######
# get the lowest y as the position for text label
 textpos = min(stats_expgroups$y0)
 
stats_expgroups %>% 
  ggplot(., aes(group = Rep)) +
  geom_boxplot(aes(x = Rep, 
                   ymin = y0, 
                   lower = y25, middle = y50, upper = y75, ymax = y100,
                   fill = expgp), 
               alpha = 0.8,
               color = "#6b6a6a",
               stat = "identity",
               size = 0.5)+
  geom_text(aes(x = Rep, y = textpos, label = str_c("n=", pepNum)))+
  geom_hline(aes(yintercept = y75ave), linetype = 1, color = "#f06e6e")+
  geom_hline(aes(yintercept = y25ave), linetype = 1, color = "#f06e6e")+
  geom_hline(aes(yintercept = y50ave), linetype = 2, color = "#f06e6e")+
  scale_fill_brewer(palette = "Set2")+
  facet_grid(. ~ expgp) +
  ylab(str_c("protein ", lb))+
  guides(fill = "none")+
  theme_bw()

ggsave(str_c("./Box plots/",la,"_Boxplot_groups_",Sys.Date(),".pdf"), plot= last_plot(), device = "pdf", width = 8, height = 6, units = "in")
ggsave(str_c("./Box plots/",la,"_Boxplot_groups_",Sys.Date(),".png"), plot= last_plot(), device = "png", width = 8, height = 6, units = "in")

remove(stats_expgroups0, stats_expgroups, gpave, stat_gp, t_gp)
dev.off()


# ====== Section 3.Visualize the isotope ratio distribution of peptides of each organism from different experiment groups =====
  

binpep <- inner_join(pepplot, OrganList, by = "Bin") 

# remove(pepplot)

# stats for each organism (keep bio replicates for t-test) #####
stats_each_rep_bin <- binpep %>%
  filter(label != "unlabeled") %>%
  group_by(Bin, Rep, diets)%>%
  summarise(minRo = min(Ratio),
            y25 = wtd.quantile(Ratio, weights = SI, probs = 0.25),
            y50 = wtd.quantile(Ratio, weights = SI, probs = 0.5),
            y75 = wtd.quantile(Ratio, weights = SI, probs = 0.75),
            maxRo = max(Ratio)) %>%
  # Note: in geom_boxplot, ymin is not the min number of y, but  
  #  "smallest observation greater than or equal to lower hinge (q1) - 1.5 * IQR"
  # where IQR ( inter-quartile range) = q3-q1
  # here instead, the following code use min of y or q1 - 1.5*IQR, whichever greater
  # similarly, ymax = max of y or q3 + 1.5*IQR, whichever smaller
  mutate(yIQR = y75 - y25) %>%
  mutate(y0 = ifelse(y25 - 1.5 * yIQR > minRo, y25 - 1.5 * yIQR, minRo),
         y100 = ifelse(y75 + 1.5 * yIQR < maxRo, y75 + 1.5 * yIQR, maxRo)) %>%
  inner_join(OrganList, by = "Bin")

write.csv(stats_each_rep_bin, str_c("./Box plots/",la, "_box_plot_data_by_Organism_Reps_",Sys.Date(),".csv"))

# Student's t-test for significance between figh fiber and high protein diet for each organism
t_or = " "
for (i in OrganList$Taxonomy) {
  # select data to test: test the medium of 3 replicates from high protein against high fiber
  stat_i <- stats_each_rep_bin %>%
    filter(Taxonomy == i) %>%
    ungroup() %>%
    select(Rep,diets,y50) %>%
  # spread the table to get the lists of high
    spread(.,diets, y50)
  
  # t-test
  t_or <- c(t_or, capture.output(cat(i), t.test(stat_i$fiber, stat_i$protein)))
}

# write t-test results
write_lines(t_or, str_c("./Box plots/",la,"_organism_t-test_",Sys.Date(),".txt"))

remove(stat_i, stats_each_rep_bin,t_or)


# stats for each organism (mix bio replicates for plotting) #####
stats_each_bin0 <- binpep %>%
  group_by(Bin, diets, label) %>%
  summarise(minRo = min(Ratio),
            y25 = wtd.quantile(Ratio, weights = SI, probs = 0.25),
            y50 = wtd.quantile(Ratio, weights = SI, probs = 0.5),
            y75 = wtd.quantile(Ratio, weights = SI, probs = 0.75),
            maxRo = max(Ratio),
            pepNum = n()) %>%
  # Note: in geom_boxplot, ymin is not the min number of y, but  
  #  "smallest observation greater than or equal to lower hinge (q1) - 1.5 * IQR"
  # where IQR ( inter-quartile range) = q3-q1
  # here instead, the following code use min of y or q1 - 1.5*IQR, whichever greater
  # similarly, ymax = max of y or q3 + 1.5*IQR, whichever smaller
  mutate(yIQR = y75 - y25) %>%
  mutate(y0 = ifelse(y25 - 1.5 * yIQR > minRo, y25 - 1.5 * yIQR, minRo),
         y100 = ifelse(y75 + 1.5 * yIQR < maxRo, y75 + 1.5 * yIQR, maxRo)) %>%
  inner_join(.,OrganList, by = "Bin")


# get ave mean of the unlabeled to add later as ref lines
bkave <- stats_each_bin0 %>%
  filter(label == "unlabeled") %>%
  select(Bin, diets, y50) %>%
  spread(diets, y50) %>%
  mutate(bkavey50 = (fiber + protein)/2) %>%
  select(Bin, bkavey50)

stats_each_bin <- inner_join(stats_each_bin0, bkave, by = "Bin") %>%
  filter(label != "unlabeled")

write.csv(stats_each_bin, str_c("./Box plots/",la, "_box_plot_data_by_Organism_",Sys.Date(),".csv"))

# get the lowest y as the position for text label
textpos = min(stats_each_bin$y0)

# plot of each bin on one graph, with unlabeled average as ref line
stats_each_bin %>% 
  ggplot(., aes(group = diets)) +
  geom_boxplot(aes(x = diets, 
                   ymin = y0, 
                   lower = y25, middle = y50, upper = y75, ymax = y100,
                   fill = Taxonomy), 
               alpha = 0.8,
               color = "#6b6a6a",
               stat = "identity", width =0.5
               )+
  geom_text(aes(x = diets, y = textpos, label = str_c("n=", pepNum)), size = 4)+
  geom_hline(aes(yintercept = bkavey50), linetype = 1, color = "#f06e6e")+
  facet_grid(. ~ Taxonomy, labeller = label_wrap_gen(width=15)) +
  ylab(str_c("Organism_Protein ", lb))+
  guides(fill = "none")+
  theme_bw()


ggsave(str_c("./Box plots/","Organism_Protein_",la,"_isotope",Sys.Date(),".pdf"), plot= last_plot(), device = "pdf", width = 24, height = 6, units = "in")
ggsave(str_c("./Box plots/","Organism_Protein_",la,"_isotope",Sys.Date(),".png"), plot= last_plot(), device = "png", width = 24, height = 6, units = "in")

remove(pepplot, stats_each_bin, OrganList)
